# Local RNA Structure, Ion Hydration Shell and the Energy Barrier for Water Exchange from the Ion Hydration Shell Determine the Mechanism of Ion Condensation on Specific RNA Sites

**DOI:** 10.1101/2022.08.15.503937

**Authors:** Antarip Halder, Sunil Kumar, Sk Habibullah, Govardhan Reddy

## Abstract

RNA folding and functioning require the binding of metal ions in specific cavities of the folded structure. This property is critical to the functioning of riboswitches that especially regulate the metal ions concentration in bacteria. However, the fundamental principles governing the specific binding of metal ions in RNA are unclear. We probed the condensation mechanism of biologically relevant alkali (Na^+^ and K^+^), alkaline earth (Mg^2+^ and Ca^2+^), and transition metals (Mn^2+^, Co^2+^, Ni^2+^ and Zn^2+^) on a part of the Ni^2+^ and Co^2+^ (NiCo) sensing riboswitch aptamer domain using computer simulations. The selected structure has multiple secondary structural elements and a single site for the specific binding of a metal ion. We show that three factors primarily determine the binding of a metal ion to an RNA site - (1) The varying structural constraints from different RNA secondary structural elements strongly influence the metal ion binding. The mode of ion binding depends on the local structure around the RNA’s ion-binding pocket. (2) The arrangement of water molecules in the ion hydration shell, and (3) the energy barrier for the ion to lose a water molecule from its hydration shell and transition from an outer to an inner shell interaction, which is primarily influenced by the metal ion charge density. These results have implications for designing biocompatible sensors using riboswitches to probe the concentration of intracellular metal ions.

## Introduction

Non-coding RNA function depends on its folding to specific compact three-dimensional structures.^1^ Intracellular metal cations condense on the RNA, and compact the polyanionic RNA chain.^2–7^ Diffusely bound metal ions on the RNA renormalize the negative backbone charge, which leads to the initial collapse of the RNA.^8^ Following that, site-bound or chelated metal ions stabilize the small electronegative pockets.^9^ Often, these site-bound ions mediate the formation of intricate tertiary interactions and stabilize the final folded state of RNA.^10^

The cations do not condense homogeneously over the polyanionic RNA backbone.^11–14^ Non-coding RNAs selectively recognize metal ions for binding in specific sites, especially in metalloriboswitches.^15,16^ Metalloriboswitches are located at the 5’-UTR of mRNAs encoding ion uptake or efflux channels in bacteria. These riboswitches are critical for maintaining ion homeostasis in bacterial cells. Upon binding by specific ions, these riboswitches fold to a structure, which signals to inhibit downstream gene expression. Till date riboswitches that can specifically sense different cations like Mg^2+^,^17^ Mn^2+^,^18,19^ Ni^2+^ and Co^2+ 20^ are discovered.

Experiments using various biophysical methods probed the principles which govern the specific recognition of metal ions by RNA.^21–28^ Computational studies complement such experiments as they can overcome the limitations imposed by the fluctuating nature of the ionic environment around RNA.^11,12,29–33^ The consensus is that hydrated metal ions can interact with the electronegative RNA atoms in two modes, viz. inner shell and outer shell (Figure 1E).^33–37^ In the inner shell mode of interaction, the ion forms a direct, coordinated covalent bond with one or more RNA atoms which requires partial dehydration of the ion, i.e., the removal of one or more water molecules from the ion’s first hydration shell.^38^ In outer shell interaction, water molecules present in the first coordination shell of a fully hydrated ion form hydrogen bonds with the RNA atoms. The mode of interaction for a specific cation is primarily determined by its charge density (*ρ*_c_).^33^ Usually, the cations with low *ρ*_c_ (like Ca^2+^) prefer the inner shell mode of interaction, whereas cations with high *ρ*_c_ (like Mg^2+^) display outer shell interactions exclusively. Computational studies have further shown that phosphate groups in RNA prefer to bind to cations with high charge density, whereas nucleobase atoms usually bind to cations with low charge density.^32^

**Figure 1:**
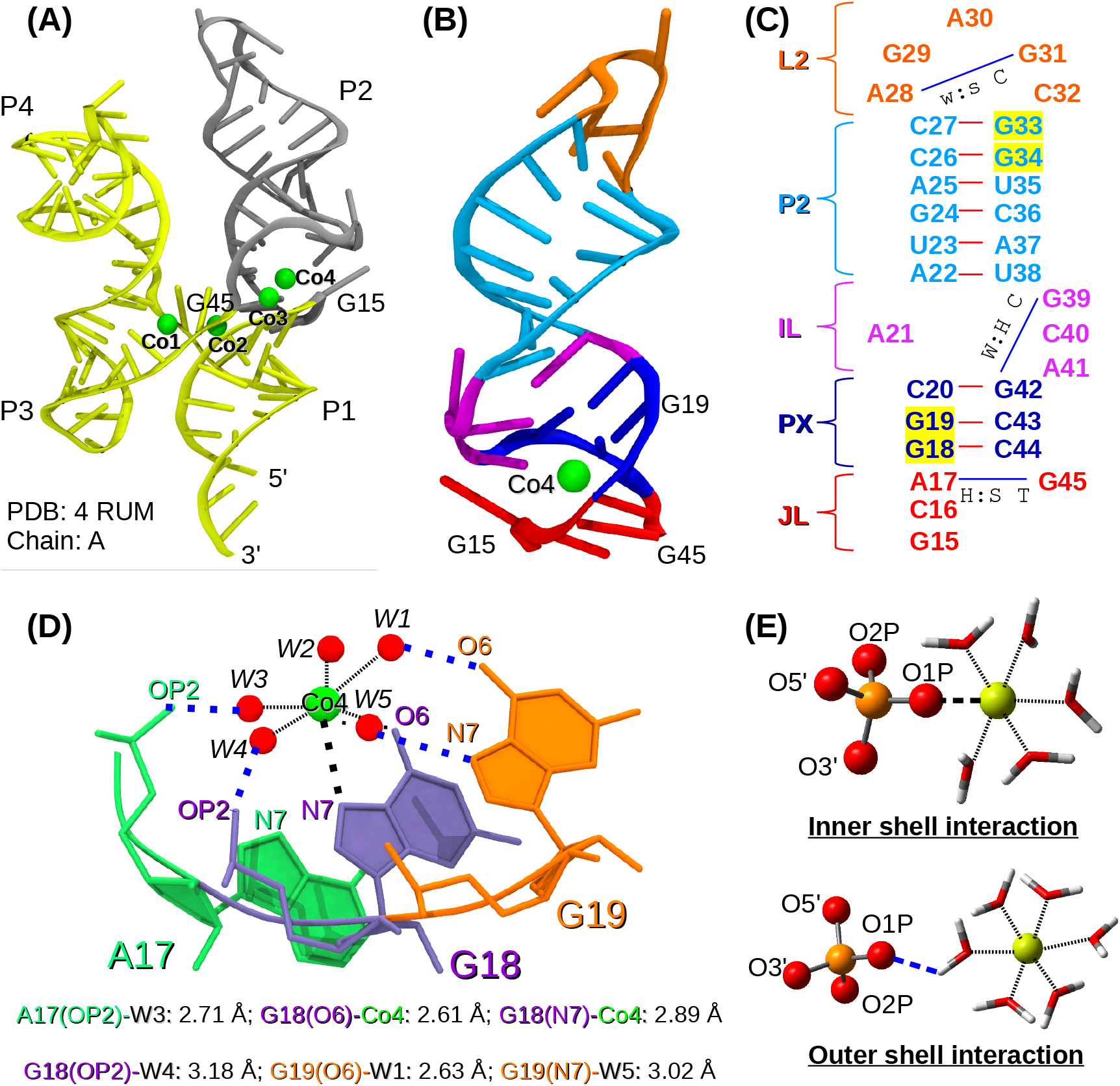
(A) Crystal structure of the aptamer domain of NiCo riboswitch^20^ in cartoon format. The bound Co^2+^ are shown as green spheres. The RNA fragment (G15-G45) selected for MD studies is highlighted in gray. (B) Three-dimensional structure of the folded G15-G45 RNA fragment and the bound Co^2+^. Secondary structural elements are highlighted (L2 loop: orange, P2 helix: light blue, IL internal loop: magenta, PX helix: deep blue, and JL junction loop: red). (C) Two-dimensional representation of the folded G15-G45 RNA fragment. Canonical (represented in red lines) and noncanonical (represented in blue lines) base pairs present in different secondary structural elements are shown. Both the PX and P2 helices have two consecutive guanine nucleotides and are highlighted in yellow. (D) Interaction of the bound Co^2+^ (Co4) and its coordinated waters with the RNA atoms. Chemical bonds between Co^2+^ and water molecules (W1-W5) present in its first coordination shell are shown as a black broken line. Water mediated interactions and direct interactions between the Co^2+^ and RNA atoms are shown in blue and black dotted lines, respectively. (E) Two modes of RNA-ion interactions are illustrated. In the inner shell mode of interaction, the RNA atom is present at the first coordination shell of the metal ion (shown as yellow sphere). In the outer shell mode of interaction, water molecules of the first coordination shell of the metal ion (shown in tube format) form hydrogen bonding interactions with RNA atoms.

The previous studies focused on the biologically prevalent alkali and alkaline earth metal ions. Despite being known for their high toxicity,^39–42^ transition metals are an integral part of many enzymes and enzyme cofactors. Therefore transition metals are indispensable for the proper functioning of various cellular processes.^43^ For example, Zn^2+^ stabilizes the structures of zinc finger proteins,^44^ Ni^2+^ constitutes the active site of hydrogenases^45^ and Co^2+^ is the central component of vitamin B_12_.^46^ Therefore, it is critical to maintaining the transition metal homeostasis in living cells to prevent both deprivation and toxicity. Bacteria utilize several classes of selective metal ion transporters to maintain metal ion homeostasis. The expression of these metal ion transporter genes is controlled by specific metal ion sensing riboswitches or metalloriboswitchs.^15^ One important example of such a metalloriboswitch is the Ni^2+^ and Co^2+^ sensing (NiCo) riboswitch, which also has biotechnological applications.^20^ It can differentiate two specific transition metals Ni^2+^ and Co^2+^ from other abundant alkali and alkaline earth metals in bacterial cells *(in vitro* ligand dissociation constant *KD* ≈ 5.6 and ≈ 12 μm for Co^2+^ and Ni^2+^, respectively).^20^ NiCo riboswitch was used in designing efficient ion sensors to detect Co^2+^/Ni^2+^ transport in *E. coli*.^47^

The NiCo riboswitch is an ideal system to probe the nature of interactions between transition metal cations and RNA. The available X-ray crystal structure^20^ of NiCo riboswitch with bound Co^2+^ ions shows that the four helices (P1, P2, P3, and P4) of the 94 nucleotides long RNA remain in a ‘twisted-H’ like shape (Figure 1A). P1 and P2 helices form one vertical side of the ‘H’ as they remain co-axially stacked. Similarly, co-axially stacked P3 and P4 helices form the other side. Twisting takes place at the four-way junction loop, which brings P1 and P3 parallel to each other. Three of the four sites where the Co^2+^ ions (Co1, Co2, and Co3) bind are located at the twisting site (Figure 1A). These RNA sites are highly specific to Co^2+^ binding, and it was argued that the binding of the three Co^2+^ ions occurs cooperatively. Compared to the three Co^2+^ ions, the fourth Co^2+^ (Co4) showed low occupancy and relatively weak anomalous density in the experiments. Also, it made fewer contacts with the conserved nucleotides (Figure 1D). Hence, the ion specificity remained ambiguous for Co4, and it was proposed that other ions may also bind at this site. Recent single-molecule experiments^48^ probed ion selectivity at the twisting site (four-way junction) of the NiCo riboswitch aptamer domain and were able to identify the folding intermediates while tracking the ion-mediated sequential twisting of the P1-P2 and P3-P4 arms.

In this work, we studied the condensation of alkali metal ions (Na^+^ and K^+^), alkaline earth metal ions (Mg^2+^ and Ca^2+^), and transition metal ions (Mn^2+^, Co^2+^, Ni^2+^ and Zn^2+^) on the P2 arm of NiCo riboswitch (residue G15 to G45, Figure 1B) using all-atom molecular dynamics simulations. Other than a few exceptions,^30,31^ ion-RNA interactions are usually studied with isolated RNA structural motifs, such as dinucleotide,^32^ helix^29^ or hairpin loop.^12^ In a large RNA structure, such isolated motifs interact with each other but under some structural constraints. Hence, the structural flexibility of the isolated motifs gets restricted. These restrictions limit the possibilities of spatial arrangements of electronegative RNA atoms. As a result, out of all the possible anionic pockets available for cation binding in isolated motifs, only a fraction is observed in large RNAs. In this study, we selected the P2 arm to overcome this limitation as it is composed of diverse secondary structure motifs, viz. four residues of the four-way junction loop (labeled as JL in Figure 1B,C), a three-base pair long helix (PX in Figure 1B,C), a six base pair long helix (P2 in Figure 1B,C), an internal loop connecting these two helices (IL in Figure 1B,C) and a penta loop capping the P2 helix (L2 in Figure 1B,C). We show that the properties that chiefly contribute to the selective ion binding to RNA are the local conformational constraints imposed by the RNA secondary structures, the rigidity of the ion water solvation shell, and the energy barrier to remove a water molecule from the primary solvation shell of the ion.

## Methods

### Unbiased MD Simulations

We have used GROMACS simulation program (version 5.1.2)^49,50^ and a modified AMBER ff14 force field^51^ along with TIP4P-EW water model^52^ to study the condensation of different metal ions on the P2 arm of the aptamer domain of NiCo riboswitch (Figure 1A,B). It is challenging for a force field to capture all the important physical properties of RNA.^53^ The modified AMBER ff14 force field was shown to sample different RNA systems reasonably well^51^ despite its drawbacks.^54^ The folded structure of the 31 nucleotide long P2 arm (G15 to G45) is taken from the X-ray crystal structure of NiCo riboswitch (PDB ID: 4RUM^20^) (Figure 1A,B). The RNA sequence is shown in Figure 1C and the phosphate group at the 5’-end is removed. The RNA is placed at the center of a cubic box of length ≈ 7 nm, and is solvated with TIP4P-EW^52^ water molecules. Periodic boundary conditions are applied in all the directions. The negative charges present on the phosphate groups along the RNA backbone are neutralized by adding 30 K^+^ ions. Additional 12 K^+^ ions are added as buffer to replicate the experimental condition. The residual charge is balanced by adding appropriate number of Cl^-^ ions. This system corresponds to a K^+^ concentration, [K^+^] ≈ 200 mM. To compare the interaction of different monovalent ions with RNA, another system is set up by adding Na^+^, instead of K^+^. This system corresponds to [Na^+^] ≈ 200 mM. To study the condensation of different divalent ions around the RNA, we prepared six additional systems with six different divalent ions (Ni^2+^, Mg^2+^, Zn^2+^, Co^2+^, Mn^2+^ and Ca^2+^) having different ionic radii (R¿) (Table 3). In these systems the concentration of the divalent ions is ≈ 60 mM, and appropriate number of Cl^-^ ions are added to maintain charge neutrality. Details of these eight systems are summarized in Table 1. Lennard-Jones (LJ) parameters for the monovalent ions are taken from the work by Joung and Cheatham.^55^ They are optimized to balance between crystal properties (e.g. lattice energy, lattice constant, etc.) and solution properties (e.g. hydration free energies). For the divalent ions, the LJ parameters developed by Li and co-workers are used.^56^ The parameters are tuned to achieve the best compromise between the hydration free energies and distance between the ion and the oxygen atoms of the water molecules in the first coordination shell.

**Table 1:** Description of the eight systems studied using simulations.

|  | Metal ion<br>concentration | No. of particles |  |  |  |  |
| --- | --- | --- | --- | --- | --- | --- |
|  |  | RNA atoms | Water | Monovalent Ion | Divalent Ion | Cl<sup>-</sup> |
| 1 | [K<sup>+</sup>] ≈ 200 mM | 1006 | 11112 | 42 | 0 | 12 |
| 2 | [Na<sup>+</sup>] ≈ 200 mM | 1006 | 11112 | 42 | 0 | 12 |
| 3 | [K<sup>+</sup>] ≈ 200 mM, [Mg<sup>2+</sup>] ≈ 60 mM | 1006 | 11076 | 42 | 12 | 36 |
| 4 | [K<sup>+</sup>] ≈ 200 mM, [Ca<sup>2+</sup>] ≈ 60 mM | 1006 | 11076 | 42 | 12 | 36 |
| 5 | [K<sup>+</sup>] ≈ 200 mM, [Mn<sup>2+</sup>] ≈ 60 mM | 1006 | 11076 | 42 | 12 | 36 |
| 6 | [K<sup>+</sup>] ≈ 200 mM, [Co<sup>2+</sup>] ≈ 60 mM | 1006 | 11076 | 42 | 12 | 36 |
| 7 | [K<sup>+</sup>] ≈ 200 mM, [Ni<sup>2+</sup>] ≈ 60 mM | 1006 | 11076 | 42 | 12 | 36 |
| 8 | [K<sup>+</sup>] ≈ 200 mM, [Zn<sup>2+</sup>] ≈ 60 mM | 1006 | 11076 | 42 | 12 | 36 |

**Table 2:** Description of different coordination number CVs used. The coordination number (*CN*) is computed using eq. 2 and 3 between the groups A and B.

| CV | To study | Group A | Group B | Remark |
| --- | --- | --- | --- | --- |
| 1 $CN_{OP-X}$ | Backbone-ion interaction | an individual ion of type $X$ present in the system | all the O1P and O2P atoms of the RNA | Higher the value of $CN_{OP-X}$, larger number of phosphate oxygen atoms are interacting with that individual ion |
| 2 $CN_X$ for G18-G19 backbone | Backbone-ion interaction | all the ions of type $X$ present in the system | sugar and phosphate oxygen atoms (O5', O3', O1P & O2P) connected to the P atoms of G18 and G19 | Value of $CN_X$ increases when any ion of type $X$ comes close to the phosphate groups of the G18 and G19 residues |
| 3 $CN_X$ for G33-G34 backbone | Backbone-ion interaction | all the ions of type $X$ present in the system | sugar and phosphate oxygen atoms (O5', O3', O1P & O2P) connected to the P atoms of G33 and G34 | Value of $CN_X$ increases when any ion of type $X$ comes close to the phosphate groups of the G33 and G34 residues |
| 4 $CN_{G18G19-X}$ | Nucleobase-ion interaction | Initially, $CN$ for each individual ion of type $X$ is computed. The individual ion of type $X$ is present in group A. The $CN$ data for all $X$ type ions is used to compute $CN_{G18G19-X}$ (see Remark) | Selected atoms of the G18 and G19 nucleobases. Selection: N1, C6, O6, C5, N7, C8 and the hydrogen atoms attached to N1 and C8 | Initially, $CN$ is computed between individual $X$-type ions (group A) and selected RNA atoms (group B). The $CN$ for all $X$ ions is used to compute the distributions involving $CN_{G18G19-X}$. When ions of type $X$ bind to the pocket composed of the G18 and G19 nucleobases, the value of $CN_{G18G19-X}$ will increase |
| 5 $CN_{U23G24-X}$ | Nucleobase-ion interaction | Initially, $CN$ for each individual ion of type $X$ is computed. The individual ion of type $X$ is present in group A. The $CN$ data for all $X$ type ions is used to compute $CN_{U23G24-X}$ (see Remark) | Selected atoms of the U23 and G24 nucleobases. Selection for U23: C4, O4, C5, C6 and the hydrogen atoms attached to C5 and C6. Selection for G24: N1, C6, O6, C5, N7, C8 and the hydrogen atoms attached to N1 and C8 | Initially, $CN$ is computed between individual $X$-type ions (group A) and selected RNA atoms (group B). The $CN$ for all $X$ ions is used to compute the distributions involving $CN_{U23G24-X}$. When ions of type $X$ bind to the pocket composed of the U23 and G24 nucleobases, the value of $CN_{U23G24-X}$ will increase |
| 6 $CN_{ions}$ for G18-G19 site | Nucleobase-ion interaction | all the ions of type $X$ present in the system | Selected atoms of the G18 and G19 nucleobases. Selection: N1, C6, O6, C5, N7, C8 and the hydrogen atoms attached to N1 and C8 | Value of $CN_{ions}$ increase when any ion of type $X$ comes close to any one of the selected atoms of G18 and G19 |
| 7 $CN_{ions}$ for U23-G24 site | Nucleobase-ion interaction | all the ions of type $X$ present in the system | Selected atoms of the U23 and G24 nucleobases. Selection for U23: C4, O4, C5, C6 and the hydrogen atoms attached to C5 and C6. Selection for G24: N1, C6, O6, C5, N7, C8 and the hydrogen atoms attached to N1 and C8 | Value of $CN_{ions}$ increase when any ion of type $X$ comes close to any one of the selected atoms of U23 and G24 |

**Table 3:** Ionic radii (*R_i_*),^77^ water-ion binding distances (*R_h_*) corresponding to the force field used,^55,56^ free energy barrier for the transition of water molecule from the first hydration shell of an ion to the second hydration shell 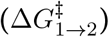 computed using metadynamics simulations, and water exchange rate constants (*k*_1_)^78–80^ for the metal ions. For Mn^2+^ and Co^2+^, the *R_i_* corresponding to their high spin states are reported since they are more stable than the low spin states. The reported *k*_1_ were estimated at 298 K in the experiments. The average lifetime (*τ*_ion_) of ion-RNA interactions is from the simulations reported in this work. For systems with a mixture of mono- and divalent ions, *τ*_ion_ of the K^+^ ion is given in the parenthesis.

| Ion | $R_i$<br>(Å) | $R_h$<br>(Å) | $\Delta G_{1 \rightarrow 2}^\ddagger$<br>(kJ/mol) | $k_1$<br>(s<sup>-1</sup>) | $\tau_{ion}$<br>(ps) |
| --- | --- | --- | --- | --- | --- |
| $Ni^{2+}$ | 0.69 | 1.91 | 36.87 | $3.2 \times 10^4$ | 568.5 (338.4) |
| $Mg^{2+}$ | 0.72 | 2.05 | 47.21 | $6.7 \times 10^5$ | 447.0 (330.6) |
| $Zn^{2+}$ | 0.74 | 1.94 | 44.40 | $3 \times 10^7$ | 491.2 (330.7) |
| $Co^{2+}$ | 0.745 | 1.97 | 48.07 | $3.2 \times 10^6$ | 554.0 (334.3) |
| $Mn^{2+}$ | 0.83 | 2.11 | 36.23 | $2.1 \times 10^7$ | 1426.7 (333.1) |
| $Ca^{2+}$ | 1.00 | 2.53 | 15.89 | $\sim 10^8$ | 2213.6 (344.9) |
| $Na^+$ | 1.02 | 2.66 | 11.20 | $\sim 10^8$ | 632.6 |
| $K^+$ | 1.58 | 2.29 | 5.69 | $\sim 10^8$ | 363.6 |

The simulation box is initially subjected to energy minimization using steepest descent minimization algorithm with a maximum force cut-off of 1000 kJ/(mol nm). The energy minimized systems are equilibrated in three steps: (i) position restraints are applied on the RNA atoms and ions using a harmonic potential with a force constant of 1000 kJ/(mol nm^2^) and only the solvent molecules are equilibrated for 2 ns in the NVT ensemble, (ii) position restraints are applied on the RNA atoms and ions and only the solvent molecules are equilibrated for 4 ns in the NPT ensemble and (iii) position restraints are applied only on the RNA atoms and both the solvent molecules and ions are equilibrated for 4 ns in the NPT ensemble. Finally, the equilibrated systems are subjected to production runs. The temperature *T* is maintained at 300 K using the stochastic velocity rescaling thermostat developed by Bussi *et al.*^57^ (with time constant *τ_t_* = 0.1 ps) and the pressure *P* is maintained at 1 atm using Parrinello-Rahman barostat^58^ (with time constant *τ_p_* = 2.0 ps). During the production run, all the bond lengths involving the hydrogen atoms are constrained using the LINCS algorithm.^59^ The simulation is advanced using the leap-frog integrator with a 2 fs time step. The long-range electrostatic interactions are computed using particle mesh Ewald summation (PME) method^60^ using 1 nm cutoff for both electrostatic and van der Waals interactions. For each system, we ran two independent trajectories of length ≈ 5 μs and ≈ 3 μs, respectively. The cumulative length of the simulation trajectories for each metal ion system is ≈ 8 μs, and for all the 8 metal ion systems is ≈ 64 μs. We saved the simulation trajectory every 5 ps for analysis. The first 250 ns of data from each trajectory are ignored and rest of the data are used for analyses.

### Well-Tempered Meta-Dynamics Simulations

We performed well-tempered metadynamics to compute the barrier height 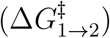 in the exchange of a single water molecule between the first and second solvation shell of hydrated metal ions. The simulations are started with a single metal ion solvated with 4138 water molecules in a cubic box of length 5 nm. All the other simulation parameters are identical to the unbiased MD simulations. Initial configurations are energy minimized, followed by equilibration. We ran each system for at least 1-3 *μ*s (depending on the system) in the NPT ensemble at 300 K using PLUMED 2.5.1^61,62^ and GROMACS 2018.6 software. We projected the two-dimensional free energy surface (FES) on two CVs: (i) distance between the metal ion and oxygen atom of water in the first solvation shell (*d*_1_) and (ii) distance between the metal ion and oxygen atom of water in second solvation shell (*d*_2_) (Figure S1A). A cut-off of 0.6 nm is used for both *d*_1_ and *d*_2_. The 1D and 2D FES for each system are constructed by sum_ hills module embedded in PLUMED and are plotted using matplotlib.^63^ In these simulations, gaussian width, initial gaussian height, deposition time, and bias factor are set to 0.01 nm, 0.25 kJ/mol, 20 ps, and 15, respectively.

### Quantifying Structural Changes

We used *eRMSD* to estimate the deviation between 2 RNA structures.^64^ Unlike the conventional root mean square deviation *(RMSD),* which uses the positions of all nucleic acid atoms to compute the deviation between 2 structures, *eRMSD* considers only the relative positions and orientations of nucleobases. *eRMSD* is shown to be more effective and less ambiguous descriptor compared to *RMSD*.^65–67^ *eRMSD* values of the RNA conformations are calculated with respect to the structure obtained after the energy minimization of the crystal structure.

For studying the structural changes occurring specifically in the backbone orientation, we calculated two dihedral angles, *η* (between 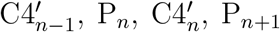) and *θ* (between 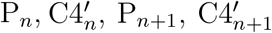) (Figure S2). Equivalent to *φ-ψ* angles in proteins, these *η* and *θ* angles are known to provide an accurate and comprehensive description of different RNA backbone conformations.^68^

### Solvent Accessible Surface Area (SASA)

The *SASA* of individual ions are calculated using the double cubic lattice method^69^ implemented in GROMACS using a water molecule radius of 0.14 nm. The normalized *SASA* values of the *i^th^* ion (*SASA^norm^*) are obtained by dividing the *SASA* of the ion with its maximum value, 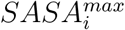. When an ion is present in the bulk solvent, its *SASA^norm^* = 1. The *SASA^norm^* decreases below 1 when the ion interacts with the RNA. The *SASA^norm^* values of individual ions are used to calculate the lifetimes of ion-RNA interactions. To avoid the transient ion-RNA contacts, only those instances were analyzed when the *SASA^norm^* value of a particular ion is below 0.8 for more than 50 ps during the simulation.

### Condensation of Metal Ions Around RNA

To quantify the condensation of positively charged metal ions onto the negatively charged RNA, we calculated the local ion concentration (*c*\*) around the (a) oxygen atoms of the phosphate groups, (b) oxygen atoms of the sugar moiety and (c) carbonyl oxygen and imino nitrogen atoms of nucleobases. Following earlier work,^11^ *c*\* in molar units is defined as,

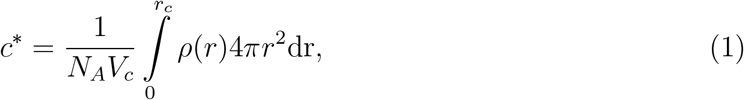

where *ρ*(*r*) is the number density of a specific type of ions at a distance *r* from a specific RNA atom, *V*_c_ is the spherical volume of radius *r*_c_, and *NA* is the Avogadro’s number. The cutoff distance, *r*_c_ is set to 0.73 nm, which is the Bjerrum length of water at *T* = 300 K.

To study the distribution of different metal ions around the RNA nucleotidess, we computed the spatial density maps using the volmap plugin of the VMD software.^70^ Initially, RNA atoms of all the conformations are aligned to the last conformation of the ≈ 5 μs trajectory. The position of metal ions in each simulation frame are moved accordingly. The alignment is performed using the Kabsch method^71^ implemented in VMD. To construct the spatial density maps, the metal ions are treated as spheres of radii equal to their respective van der Waals radii. A grid of size 0.01 nm^3^ is constructed to divide the space around the RNA nucleotides. A grid point is considered to be occupied and assigned a value equal to 1, if the sphere corresponding to a specific metal ion lies on the grid. Otherwise it takes a value equal to 0. The value at a grid point is averaged over all frames of the trajectory. VMD is used for structural superposition^72^ and rendering of images.^73^

### Collective Variable (CV) to Distinguish Metal Ion Interaction With the RNA Atoms

A CV named coordination number (*CN*) is used to quantify the proximity of metal ions to the backbone, nucleobases and specific binding pockets of the RNA. The COORDINATION keyword of PLUMED package^61,62^ (version 2.5.2) is used to calculate CN between two groups of atoms (say group A and group B) over a given MD trajectory. At each time step, we calculated the switching function *S_ij_* given by

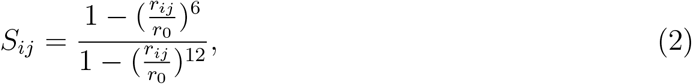

where *r*_ij_ is the inter-atomic distances between the atoms of group A (indexed with *i*) and atoms of group B (indexed with *j*), and *r*_0_ is the cut-off distance and is taken to be 0.63 nm to capture both the inner shell and outer shell interactions between the ion and the RNA (Figure 1). Finally, CN is calculated as

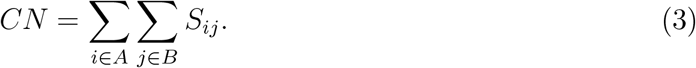

To measure the proximity of phosphate oxygen atoms (O1P/O2P) to the individual ions of type *X*, the *CN* was calculated between (a) an individual ion of type *X* present in the system and (b) all the O1P and O2P atoms of the RNA. We represent this CV as *CN*_OP-*X*_. For example, if *X* = Ni^2+^, then the CV will be *CN*_OP-*Ni*_. Higher the value of *CN_OP-X_*, larger number of phosphate oxygen atoms are interacting with that individual ion. For each time step in the MD trajectory, this calculation is repeated for every ion of type X present in the system.

To compute *CN*_G18G19–*X*_, *CN* was initially computed for all the individual ions of type *X*. In computing *CN*, group A in eq. 3 consists of an individual ion of type *X*, and group B consists of selected atoms of the G18 and G19 nucleobases. These selected atoms are N1, C6, O6, C5, N7, C8 and the hydrogen atoms attached to N1 and C8. The CN of all the ions of type *X* are combined to compute *CN*_G18G19–X_. When an ion of type *X* binds to the pocket composed of the G18 and G19 nucleobases, the value of *CN*_G18G19–*X*_ will increase.

Similarly, we calculated *CN*_U23G24–*X*_, whose value will increase when an ion of type *X* binds to the pocket composed of the nucleobases of the U23 and G24 residues. To compute CN required for the computation of *CN*_U23G24–*X*_, the selected group B atoms of U23 are C4, O4, C5, C6 and the hydrogen atoms attached to C5 and C6. The same for G24 are N1, C6, O6, C5, N7, C8 and the hydrogen atoms attached to N1 and C8.

### Joint Probability Distributions

The probability distribution (PD) corresponding to ion binding in negative logarithmic scale is calculated using *P*_L_(**s**) = – ln(*P*(**s**)), where *P*(**s**) is the probability distribution of the set of CVs **s**. We estimated *P*(**s**) by histogramming the values of the CVs **s** computed from the simulation data. The PDs are plotted using matplotlib tool.^63^ To probe the interaction between ions of type *X* and the RNA backbone, we computed the joint probability distribution (JPD) of two CVs in negative logarithmic scale (i) *CN*_OP-*X*_ and (ii) minimum distance between an individual ion *X* and any of the phosphate oxygen atoms of the RNA (min(*d*_OP-*X*_)). *CN*_OP-*X*_ does not discriminate between the inner shell and outer shell coordination between ion and phosphate oxygen atoms. Whereas, min(*d*_OP-*X*_) can differentiate between the two modes of interaction. Similarly to probe the interaction between ions of type *X* and binding pockets composed of G18 and G19 nucleobase atoms, we computed JPD of two CVs (i) *CN*_G18G19-X_ and (ii) minimum distance between an individual ion *X* and any of the electronegative nucleobase atoms (min(*d*_Base-X_))· Similarly, we computed JPD of (i) *CN*_U23G24-*X*_ and (ii) min(*d*_Base-*X*_).

## Results

### Binding Mode Determines the Binding Lifetime of Divalent Ions to RNA

To probe the effect of different ionic conditions on the structure of P2 arm, we quantified the structural changes by computing *eRMSD* for the whole RNA and its individual secondary structural elements (JL, IL, L2, PX, and P2) (Figure 1B, 1C, S3 and S4). A minor increase in *eRMSD* (≈ 1) is observed in all the eight systems studied, suggesting that the P2 arm retains its native-like structure for different ionic conditions (Figure S3 and S4). The observed increase in *eRMSD* for the whole RNA is primarily due to the fluctuations in the loop regions (Figure S3, S4, and S5). The stable P2 arm structure with sequence diversity and multiple secondary structural elements is ideal for studying the role of local RNA structural heterogeneity on the ion-RNA interactions and ion binding modes.

Previous studies reported that charge density (*ρ*_c_) values of cations determine the nature of ion-RNA interactions, at least for the alkaline earth metals.^11,32^ For example, low *ρ*_c_ ions like Ca^2+^ can either interact with RNA through the outer shell mode (water-mediated ion-RNA interaction) or lose a water molecule from its first solvation shell to interact with the RNA through inner shell mode (direct ion-RNA interaction) (Figure 1E). Whereas high *ρ*_c_ ions like Mg^2+^ interact with RNA only through the outer shell mode.

When metal ions interact with RNA, either by binding at specific-sites or being diffused along the RNA contour, their respective SASA decreases significantly. We observed that the SASA of individual metal ions increased and decreased multiple times during the simulation, indicating multiple ion binding and unbinding events (Figure S6 and S7). A simple argument regarding ion binding is that the average ion binding lifetime (*τ*_ion_) increases with the increase in *ρ*_c_.^32,74^ However, it holds only for the monovalent alkali metals (K^+^ and Na^+^) studied here. *ρ*_c_ and *τ*_ion_ of Na^+^ are higher than those of K^+^ (Table 3). The divalent alkaline earth (Mg^2+^ and Ca^2+^) and transition metals (Mn^2+^, Co^2+^, Zn^2+^ and Ni^2+^) do not follow this trend. Rather, the divalent ions with smaller *ρ*_c_ have larger *τ*_ion_, and Ni^2+^ is the only exception to this trend among the ions we studied (Table 3). Despite Ni^2+^ having the largest *ρ*_c_, *τ*_ion_ of Ni^2+^ is only moderately larger than that of Mg^2+^, Zn^2+^ and Co^2+^ (Table 3).

The additional factor that determines the *τ*_ion_ for a particular ion is the presence of other competing cations in the same system. The *τ*_ion_ of K^+^ decreased in presence of divalent ions in the solution (Table 3). The water-ion binding distance (R_*h*_) for K^+^ is higher than any of the divalent ions (Table 3). The hydrated divalent ions have a larger charge to volume ratio compared to hydrated K^+^. As a result, in a mixture of mono- and divalent ions, divalent ions are preferred over the monovalent ions for condensation around RNA.^11,12^

For divalent ions, our observations align with the claim that *τ*_ion_ depends on the mode of ion binding.^33^ The inner shell coordination is thermodynamically more stable than the outer shell coordination for ion-RNA interaction.^75,76^ Hence, *τ*_ion_ of ions binding to RNA atoms through inner shell mode is larger than that of ions binding through outer shell mode. The *τ*_ion_ for Mn^2+^ and Ca^2+^ as obtained from our simulations are significantly higher compared to all other divalent ions considered in this work (Ni^2+^, Mg^2+^, Zn^2+^ and Co^2+^)(Table 3). Generally, ions with low *ρ*_c_ can form inner shell coordination with RNA as they can easily lose the water molecules from their first solvation shell.^33^ The alkaline earth metal Ca^2+^ and the transition metal Mn^2+^ have smaller *ρ*_c_ than other divalent ions. They are also associated with faster water exchange rates (*k*_1_) (Table 3). As a result, Mn^2+^ and Ca^2+^ can get partially dehydrated and interact with the RNA atoms through inner shell mode. We further probed the ion binding around multiple RNA sites to understand the underlying factors contributing to the preferential condensation of ions on some specific RNA sites (Figure 2A,B).

**Figure 2:**
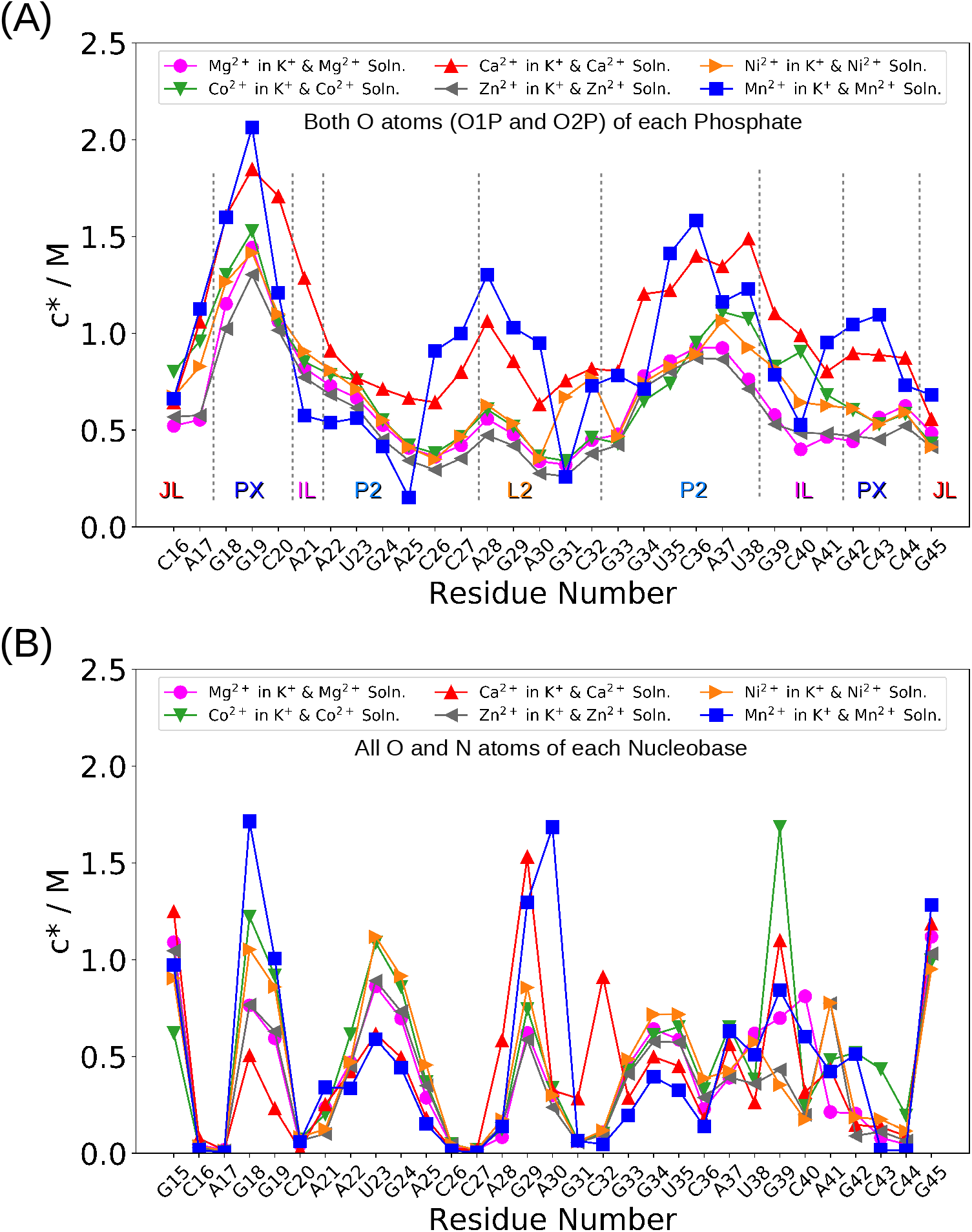
Local concentration (*c*\*) of the divalent ions around (A) phosphate oxygen atoms (both O1P and O2P) and (B) the nucleobase atoms (only N and O) are shown for Ni^2+^, Mg^2+^, Zn^2+^, Co^2+^, Mn^2+^ and Ca^2+^. Ions bind preferentially to both phosphates and nucleobases of the nucleotides G18-G19, and only to the nucleobases of nucleotides U23-G24. The ions show preference for the nucleobase of nucleotide G29 located in the loop region. Ca^2+^, Mn^2+^, Mg^2+^, and Co^2+^ exhibit preferential condensation around the solvent exposed nucleobase of nucleotide G39.

### Backbone Orientation Determines the Extent of Ion Condensation at Similar RNA Sites

To probe the cation condensation around the RNA, we computed the local concentration (*c*\*) of the mono- and divalent ions around the phosphate and nucleobase sites. We used *r*_c_ = 0.73 nm for phosphate oxygen atoms (O1P and O2P) and *r*_c_ = 0.5 nm for nucleobase N and O atoms while computing the *c*\* (eq. 1). The ions with high *ρ*_c_ condense at specific phosphate sites, whereas ions with low *ρ*_c_ diffuse over the whole RNA (Figure 2A,B and S8). In addition to the backbone (Figure 2A), there are two specific sites on the RNA where the ions condense around the nucleobase sites (Figure 2B). These nucleobase sites are located on the major grooves of the PX (around G18 and G19), P2 (around U23 and G24) helices, and L2 loop (G29). Ions show comparatively less condensation at the nucleobase sites G34 and U35 (Figure 2B). Ca^2+^, Mn^2+^, Mg^2+^, and Co^2+^ ions also condense around the solvent exposed nucleobase of nucleotide G39. The high *c*^*^ value for all the ions at the nucleobases G15 and G45 is because these nucleobases are located at the terminal with easy access to the cations. The crystal structure reports the bound Co^2+^ (Co4 ion) at the Hoogsteen edges of the nucleobases of G18 and G19 (Figure 1B,D). No bound Co^2+^ is reported around U23 and G24 in the crystal structure.^20^

The value of *c*\* around the phosphate (O1P and O2P atoms) and sugar groups (O2’, O3’, O4’ and O5’ atoms) reveals ion condensation around the RNA backbone. The degree of ion condensation around O1P, O2P, O3’ and O5’ is higher than that around other O atoms (Figure S9A,B and S10B,D). These are the atoms that are involved in the formation of the phosphodiester linkage between two consecutive nucleotides. In general, *c*\* of Ca^2+^ and Mn^2+^ remain higher than the ions with high *ρ*_c_ (Ni^2+^, Mg^2+^, Zn^2+^ and Co^2+^) due to their inner shell coordination with RNA as observed on the simulation time scale (see next section). Analyses of the variation in SASA values also showed that ion-RNA interactions have longer lifetimes for Mn^2+^ and Ca^2+^ (Figure S6E,F, S7E,F, and Table 3). Interestingly, *c*\* around phosphate groups for all the ions is maximum around the 5^’^ end of the PX helix, especially around G18 and G19 (Figure 2A). The other helix P2 also has two consecutive guanine residues G33 and G34 (Figure 1C). But, *c*\* for all the ions are relatively lower around G33-G34 as compared to the G18-G19 nucleotides (Figure 2A). To understand the difference in the ion condensation around similar set of nucleotides, we probed the backbone conformations of nucleotides G18-G19 (PX helix), and nucleotides G33-G34 (P2 helix) (Figure 1B,C).

Backbone conformations are quantified by the dihedral angles *η* and *θ*^68^ (see Methods). For the consecutive residues (say for G18-G19) we calculated *η* and *θ* for both the residues (i.e. *η*18, θ18 and *η*19, *θ*19). The distribution of *η* and *θ* is projected in the form of a scatter plot (Figure 3, S11 and S12). The backbone at nucleotides G18-G19 sample only a limited region in the *η-θ* space that is centered around |*η*| ≈ 150° and |*θ*| ≈ 150° (Figure 3A,B S11A,B and S12A,B). Whereas, the backbone at nucleotides G33-G34 is more flexible and can adopt a vast number of other conformations that have lower values of |*η*| and |*θ*| (Figure 3C,D S11C,D and S12C,D). The electrostatic potential around the backbone of a nucleotide *n*, which contributes to ion binding is determined by the spatial organization of the electronegative O atoms attached to its P atom (i.e. O1P(*n*), O2P(*n*), O3’(*n* – 1) and O5’(*n*)).

**Figure 3:**
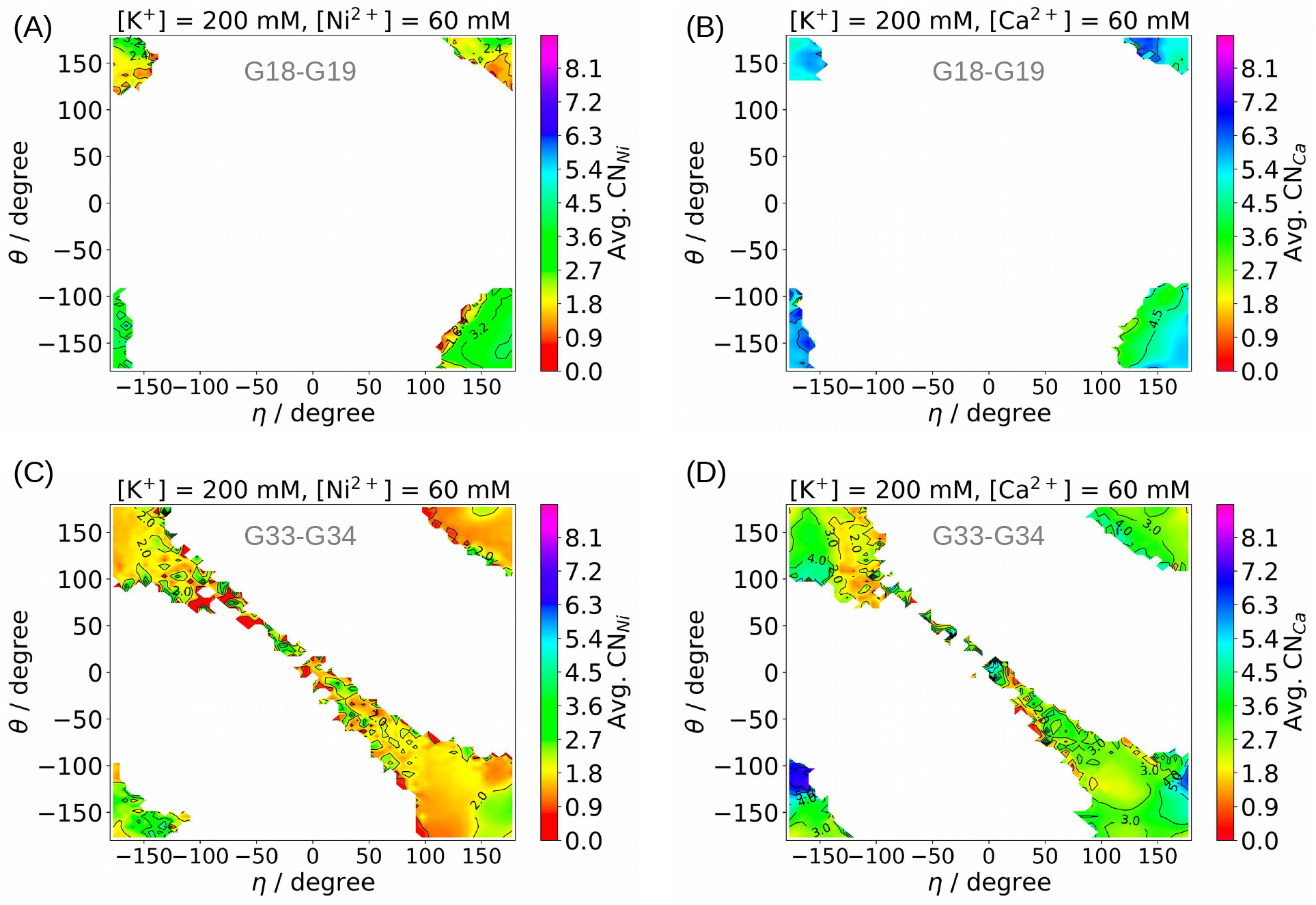
Scatter plots of the dihedral angles *η* and *θ* to map the backbone conformations adopted by the G18 and G19 nucleotides for systems (A) Ni^2+^ and (B) Ca^2+^. Each point is colored on the basis of the average values of the *CN*_X_, where *X* = Ni^2+^, Ca^2+^, etc. For a nucleotide *n, CN*_X_ is the CN measured between (i) four O atoms viz. 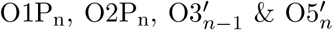 and (ii) all the ions of type *X* present in the system. The average value of *CN*_X_ for a nucleotide n increases when ions of type *X* bind to its backbone. The results obtained for G18 and G19 nucleotides are compared with the distribution of the G33 and G34 backbone orientations for (C) Ni^2+^ and (D) Ca^2+^.

We colored each point in the scatter plot according to the 〈*CN*_X_〉 (Table 2) (Figure 3, S11 and S12). Higher the value of 〈*CN*_X_〉, the greater the frequency of ion binding at the backbone (see Methods and Table 2). In all the systems, 〈*CN*_X_〉 is high at majority of the conformations adopted by G18-G19, i.e. at the regions centered around |*η*| ≈ 150° and |*θ*| ≈ 150^°^. Whereas the conformations that are exclusive to G33-G34 are almost always associated with a low 〈*CN*_X_〉. Hence, only a specific set of backbone orientations can provide the complementary electronegative environment required for the cation binding. The backbone of G18-G19 nucleotides is constrained around those specific orientations which facilitate cation binding. Such structural conformations are absent for the G33-G34 backbones. We hypothesize that even in the absence of tertiary interactions, the secondary structures of RNA are sufficient to impose constraints on certain conformations of the polyanionic backbone that are favorable for metal ion binding. This explains why the metal ions condense around their specific backbone binding site, even when the RNA is in its unfolded state.^11–14,81^ We further analyzed the interaction between individual ions and RNA atoms (phosphate O atoms and nucleobase atoms), which revealed the difference in the interaction mechanism for alkaline earth and transition metals.

### Ions with Low Energy Barrier 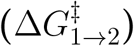 to Lose a Water Molecule From Their First Solvation Shell Switch Between the Inner and Outer Shell Binding Modes

We computed the joint probability distribution (JPD) to identify various states in metal ion binding to the RNA backbone using the two CVs: (i) coordination number (CN) (see eq. 2 and 3) between an individual ion and all the phosphate O atoms (*CN*_OP-*X*_, where *X* = Ni^2+^, Mg^2+^, etc.), and (ii) the minimum distance between that specific ion and any of the phosphate O atoms (min(*d*_OP-*X*_), where *X* = Ni^2+^, Mg^2+^, etc.) (see Methods and Table 2). The higher value of *CN*_op-*X*_ indicates that a larger number of phosphate O atoms interact with the individual ion. The CV min(*d*_OP-*X*_) can distinguish between inner and outer shell coordination. The region in the JPD centered at min(*d*_OP-*X*_) ≈ 0.4 nm corresponds to the outer shell coordination, where the metal ion interacts with O1P/O2P via its coordinated water molecules (Figure 4). The region in the JPD centered at min(*d*_OP-*X*_) ≈ 0.2 nm corresponds to the inner shell coordination, where the metal ion interacts directly with O1P/O2P.

**Figure 4:**
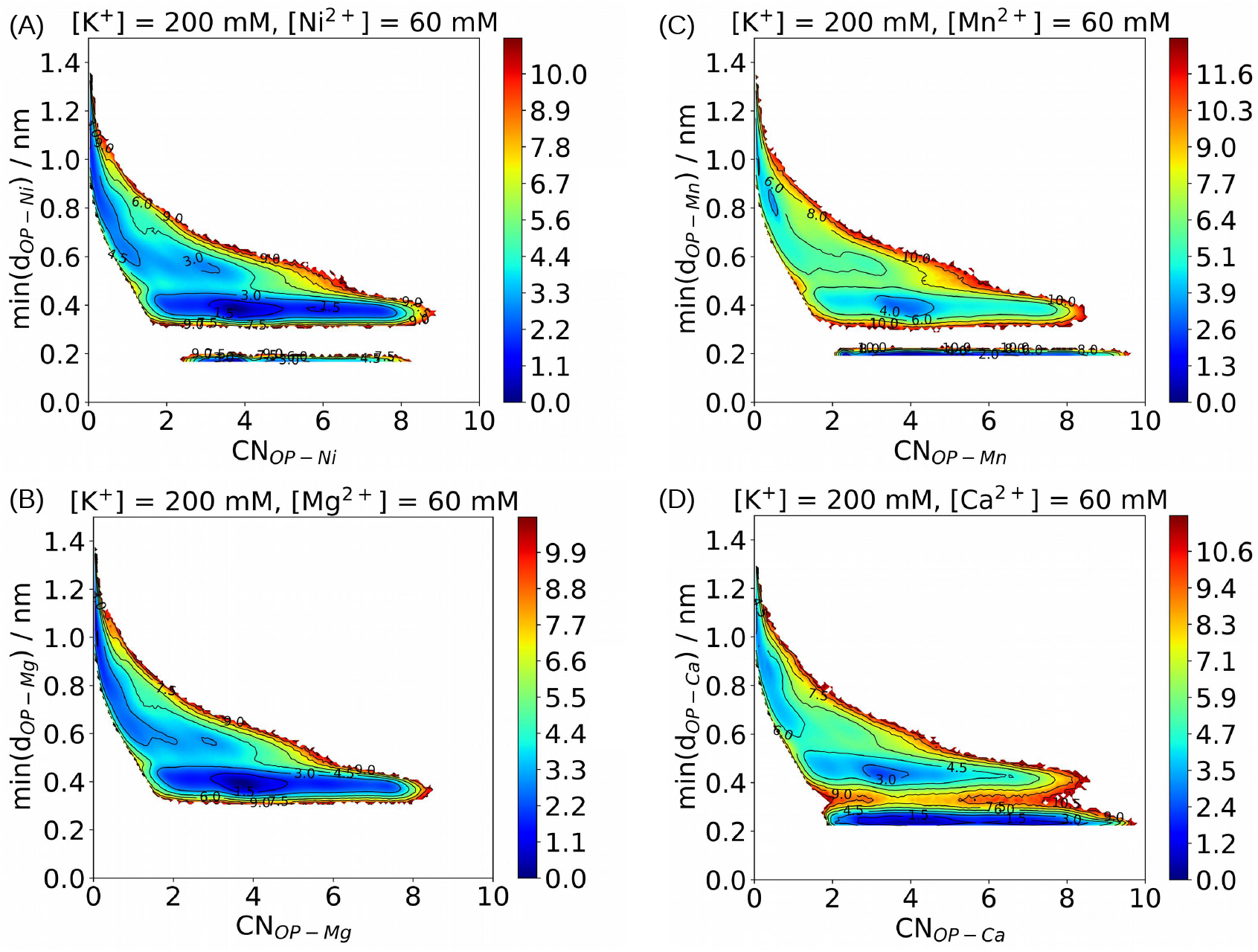
JPD is plotted in negative logarithmic scale for the binding of ions with the RNA backbone for systems (A) Ni^2+^ (transition metal with high *ρ*_c_) (B) Mg^2+^ (alkaline earth metal with high *ρ*_c_) (C) Mn^2+^ (transition metal with low *ρ*_c_) and (D) Ca^2+^ (alkaline earth metal with low *ρ*_c_). The JPD is plotted for *CN*_OP-*X*_ and min(*d*_OP-*X*_). *CN*_OP-*X*_ is the coordination number between an individual ion of type X and all the phosphate O atoms. min(d_OP-X_) is the minimum distance between that particular ion of type X and any of the phosphate O atoms. Similar analyses for the other ions are shown in Figure S13.

Ni^2+^ and Mg^2+^ are the transition and alkaline earth metal ions, respectively, with the high *ρ*_c_ value. Ni^2+^ can form both inner and outer shell coordination, although the outer shell coordination remains the preferred mode of interaction with phosphate O1P/O2P atoms (Figure 4A). Mg^2+^ only forms outer shell coordination with the backbone^11^ (Figure 4B). The other transition metals (Zn^2+^ and Co^2+^) with *ρ*_c_ higher than Mn^2+^ formed only outer shell coordination with the RNA backbone on the simulation time scale (Figure S13A,B). Ca^2+^ and Mn^2+^ with low *ρ*_c_ form both inner shell and outer shell coordination with the phosphate O1P/O2P atoms (Figure 4C,D). Since the inner shell ion-RNA interaction is comparatively more stable, we observed a longer ion-RNA binding lifetime for Ca^2+^ and Mn^2+^ compared to other divalent ions (Table 3, Figure S6E,F and S7E,F). The alkali metal ions, Na^+^ (Figure S13C) and K^+^ (Figure S13D), also populated conformations where both inner and outer shell coordination with RNA are observed, but as *ρ*_c_ of Na^+^ and K^+^ are even lower than that of Ca^2+^, both alkali ions show comparatively low binding lifetime with RNA.

The area between the regions corresponding to the inner and outer shell coordination in JPD has non-zero probability in the case of Ca^2+^, Na^+^, and K^+^ (Figure 4D and S13C,D). For Ca^2+^, we observed multiple transitions between the outer and inner shell coordination modes of binding (Figure S16). In contrast, for Mn^2+^ and Ni^2+^, the regions corresponding to inner and outer shell coordination in JPD are disjoint (Figure 4A,B). For Mn^2+^, we observed multiple events, where the ion switches from outer to inner shell binding. We did not observe inner shell to outer shell binding transitions on the simulation timescale (Figure S15). Changes in the min(*d*_OP-*X*_) values show that Ni^2+^ and Mn^2+^ switch from outer to inner shell coordination while interacting with O1P/O2P (Figure S14 and S15). But once the inner shell coordination is formed, they do not switch back to outer shell coordination on the simulation time scale. However, for Ni^2+^ we observed an anomalous behavior, where despite its high *ρ*_c_ and low experimental *k*_1_ values (Table 3) it switches from outer to inner shell coordination. We found that the switch occurred only once in the entire simulation (panel Ni3 in Figure S14A).

To understand the anomalous behavior of Ni^2+^ with respect to the *ρ*_c_ and *k*_1_ values, we performed metadynamics simulations of metal ions in water (see Methods). We used these simulations to compute the barrier height for the transition of a water molecule from the first to the second hydration shell of a metal ion 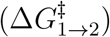 (Table 3). Low 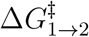 values for an ion indicate the facile transition of a water molecule from the first hydration shell, which facilitates the ion’s interaction with the RNA directly through the inner shell coordination. We found that the 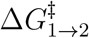 has the following order for the divalent cations, Co^2+^ > Mg^2+^ > Zn^2+^ > Ni^2+^ ≈ Mn^2+^ > Ca^2+^ (Table 3). The high values of 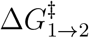 for Co^2+^, Mg^2+^, and Zn^2+^ ions allow them to interact with phosphate O1P/O2P atoms only through the thermodynamically less stable outer shell coordination leading to lower RNA binding lifetime on the simulation timescale (Table 3). The comparatively low value of the 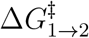 enables Ca^2+^ ion to shed water molecule from the first hydration shell and interact with phosphate O atoms (O1P/O2P) through inner shell coordination leading to higher RNA binding lifetime (Table 3). The 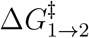 value for Ni^2+^, which is comparable to Mn^2+^ explains why it forms the inner sphere coordination despite its high *ρ*_c_ value. The computed 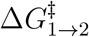 values are not in correlation with the experimentally measured *k*_1_ values (Table 3) probably due to the ion force field used in the simulations. The 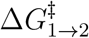 for water exchange between the hydration shells of metal ions is known to be sensitive to the metal ion parameters, and the water model used in the simulations.^82–84^ However, this analysis indicates that 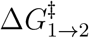 is a better indicator of whether an ion can interact through inner or outer shell coordination compared to *ρ*_c_.

### RNA Local Environment Determines the Mechanism of Ion-Nucleobase Interactions

We observed high ion condensation around two RNA nucleobase sites: (i) the major groove between G18 and G19 and (ii) the major groove between U23 and G24 (Figure 2B). We probed the nature of ion binding at these two pockets (G18-G19 and U23-G24) by computing the JPD for binding of metal ions with the nucleobase atoms (Figure 5 and S17). The JPD is computed for the two CVs: (i) *CN*_G18G19-*X*_ and *CN*_U23G24-*X*_, where *X* = Ni^2+^, Mg^2+^, etc., and (ii) the minimum distance between the *X* ions and any of the imino nitrogen atoms (N_*b*_) and carbonyl oxygen atoms (O_*b*_) of the nucleobases, min(*d*_Base-*X*_) (see Methods for details and Table 2).

**Figure 5:**
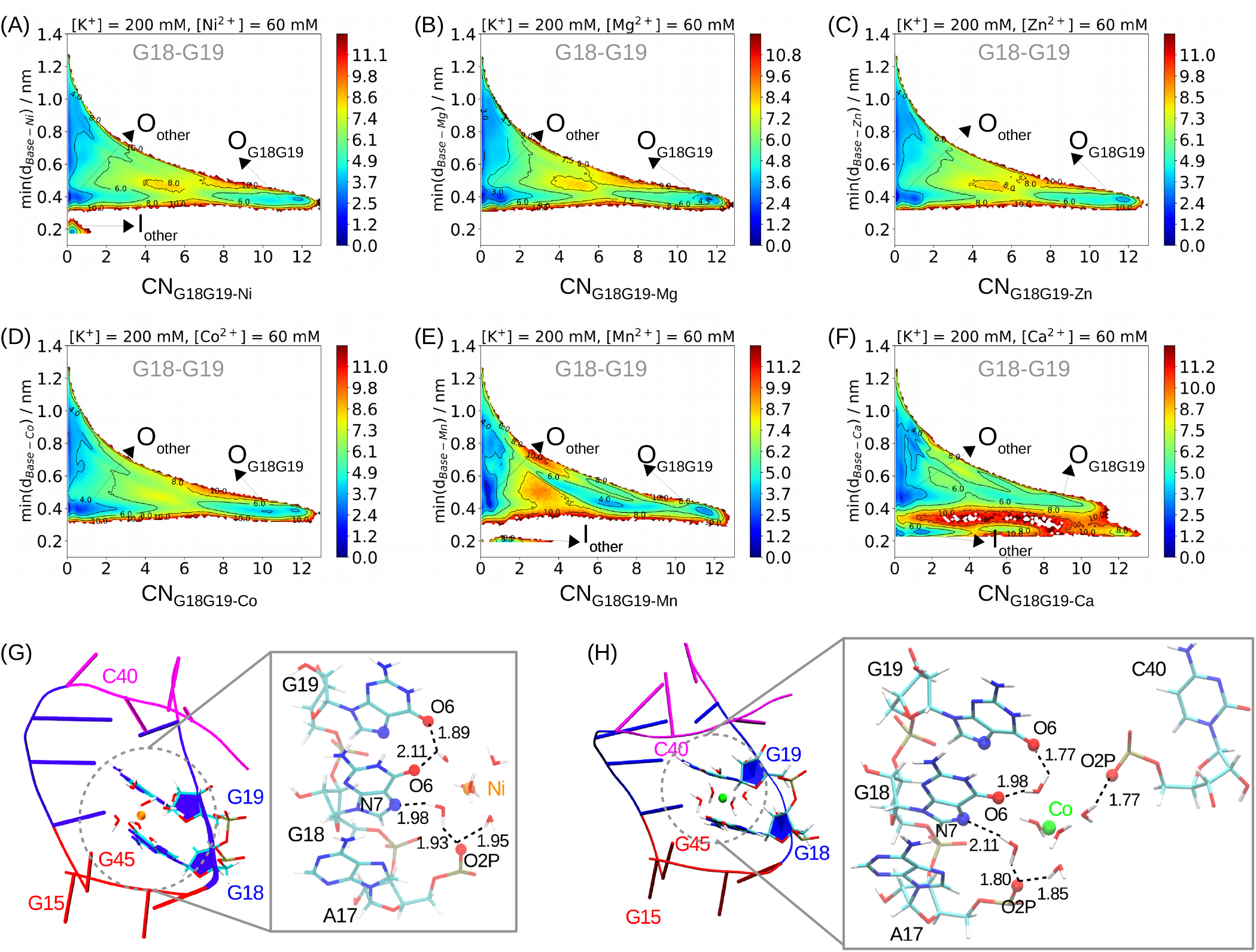
JPD in negative logarithmic scale is plotted for the binding of ions with the Hoogsteen edges of the G18 and G19 nucleotides for systems: (A) Ni^2+^, (B) Mg^2+^, (C) Zn^2+^, (D) Co^2+^, (E) Mn^2+^ and (F) Ca^2+^. The JPD is plotted for CN_G18G19-X_ and min(d_Base-*-X*_)· *CN*_G18G19-*X*_ is the coordination number between all the divalent ions of type X and selected atoms of the Hoogsteen edges of the nucleobases of G18 and G19 (see Table 2). min(*d*_Base-*X*_ is the minimum distance between all divalent ions of type X and any of the carbonyl oxygen or imino nitrogen nucleobase atoms present in the system. The regions of the JPD corresponding to the outer shell interaction with the Hoogsteen edge of G18-G19 (O_G18G19_), outer shell interaction with the nucleobase atoms other than those of G18-G19 (O_other_), inner shell interaction with the Hoogsteen edge of G18-G19 (I_G18G19_) and inner shell interaction with the nucleobase atoms other than those of G18-G19 (I_other_) are labeled. (G) Interaction between the hydrated Ni^2+^ and the binding site composed of G18 and G19 nucleobase atoms. (H) Interaction between the hydrated Co^2+^ and the binding site composed of G18 and G19 nucleobase atoms. Hydrogen bonds between ion-coordinated water and RNA atoms are represented in black broken lines with the hydrogen-acceptor distances mentioned in Å. The binding of other ions at the G18-G19 binding pocket is illustrated in Figure S19.

Inner shell and outer shell coordination between ions and N_*b*_/O_*b*_ occur at min(*d*_Base-*X*_) ≈ 0.2 nm and ≈ 0.4 nm, respectively. Similar to the ion-backbone interactions, Ni^2+^, Mn^2+^ and Ca^2+^ form both inner and outer shell coordination with the nucleobases (Figure 5A,E,F and S17A,E,F). However, the G18-G19 and U23-G24 binding pockets display specific trends. The regions in JPD for inner shell interactions (min(*d*_Base-*X*_) ≈ 0.2 nm) are observed only at lower values of CN_G18G19-*X*_. The binding of a metal ion to the pocket composed of G18 and G19 bases corresponds to a high value of CN_G18G19-*X*_ (> 10). Except for Ca^2+^, none of the other ions have a non-zero probability at *CN*_G18G19-*X*_ > 10 and min(*d*_Base-*X*_) ≈ 0.2 nm in JPD (Figure 5). Ion binding at nucleobase sites of G18-G19 pocket takes place only through outer shell interaction for divalent ions other than Ca^2+^. The JPD shows that in addition to Ca^2+^, Ni^2+^ also binds to the U23-G24 pocket using both inner and outer shell interaction (Figure S17A,F). Hence, *ρ*_c_ for metal ions is not the sole criterion to determine the nature of condensation around nucleobase atoms. The local environment around an individual site is also critical.

From the above analyses it is not clear whether these pockets are selective towards any specific ion. At G18-G19, a wide region corresponding to the ion bound states is observed for Ni^2+^, Mg^2+^, Zn^2+^ and Co^2+^ (Figure 5A,B,C,D). It is centered around min(*d*_Base-*X*_) ≈ 0.4 nm and *CN*_G18G19-*X*_ ≈ 11.5. The same is also true for U23-G24 (Figure S17). However, at the Co4 binding site of the crystal structure (Figure 1A,B), i.e. at around the Hoogsteen edges of G18 and G19,^20^ Co^2+^ shows a higher degree of condensation than other divalent ions even in the simulations. Around N7 and O6 atoms of G18 and G19, *c*\* of Co^2+^ is higher than other ions with high *ρ*_c_ (i.e. Ni^2+^, Zn^2+^ and Mg^2+^) (Figure S20A,B). This trend is not followed at U23 and G24. Except Mn^2+^, *c*\* of all other ions are nearly the same around O4 of U23 (Figure S22C,D) and N7 and O6 of G24 (Figure S20A,B). Therefore, G18-G19 pocket is to some extent selective towards Co^2+^, whereas the U23-G24 pocket is neutral to all the ions.

## Conclusion

How cations preferentially condense at specific RNA sites is a fundamental question in RNA folding and function. Using computer simulations, we probed the principles that dictate the site-specific binding of biologically relevant alkali, alkaline earth, and transitional metal ions to RNA using the P2 arm of the NiCo riboswitch aptamer domain.

We provide evidence that the following RNA-ion properties contribute to the specific binding of an ion on RNA: (1) The local RNA secondary structure plays a vital role in ion binding. Even though the local RNA sequence is identical, different secondary structures lead to variation in the electrostatic potential surface, which determines ion binding. Hence, specific secondary structure formation during RNA folding can guide the metal ions to bind at certain sites even when the tertiary interactions are absent.^11–14^ (2) To bind in specific RNA pockets, ions may have to transition from an outer to an inner shell binding through the loss of water molecules from their solvation shell to better fit into the binding pocket and strengthen their interaction with the RNA. The free energy barrier to remove a water molecule from the ions solvation shell is a better indicator for this transition than the ion’s charge density. (3) The ion solvation shell’s rigidity determines the ion’s interaction with the RNA atoms in the binding pocket.^12^ These principles can be utilized in the rational design of RNA sequences, which selectively bind to toxic metal ions present in low concentrations,^47^ that can be used in developing sensors of toxic ions.

## Acknowledgement

This work was supported in part by a grant from the National Supercomputing Mission (NSM) through grant MeitY/R&D/HPC/2(1)/2014. A.H. acknowledges the financial support from the DBT-RA Program in Biotechnology and Life Sciences. S.K. acknowledges research fellowship from the Indian Institute of Science, Bengaluru. S.H. acknowledges research fellowship from the prime minister’s research fellows (PMRF) scheme. The computations are performed using the TUE and Cray XC40 clusters at IISc.

## Notes

### Competing Interest Statement

The authors have declared no competing interest.

### Summary of Updates

Author List

