## Supplementary Figures and Tables for "Local RNA Structure, Ion Hydration Shell and the Energy Barrier for Water Exchange from the Ion Hydration Shell Determine the Mechanism of Ion Condensation on Specific RNA Sites"

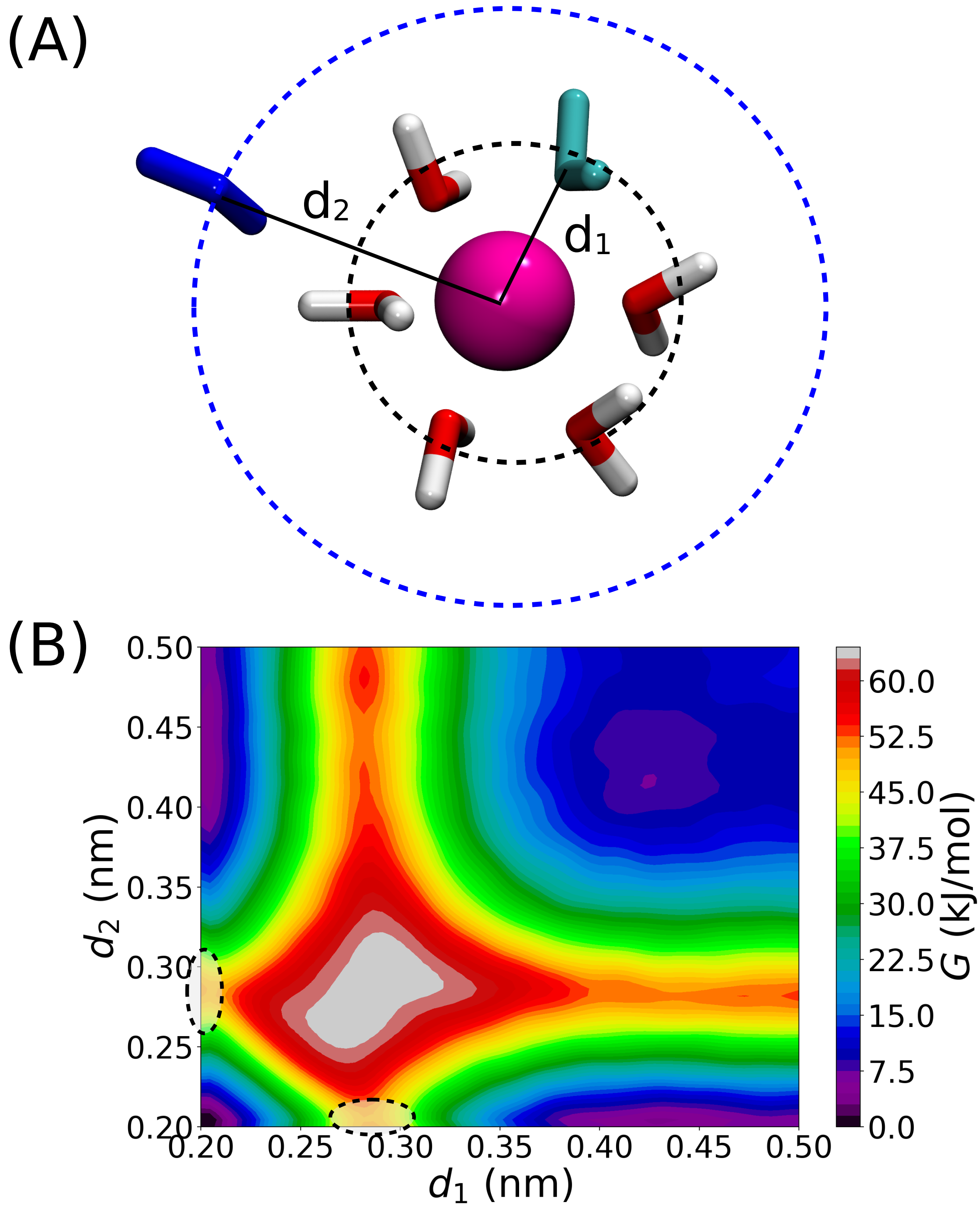


Figure S1: (A) Schematic explaining the 2 CVs chosen for the metadynamics (see Methods). (B) 2D-FES for the water exchange in hydrated $Mg^{2+}$ ion. The values reported for $\Delta G^{\ddagger}_{1\rightarrow 2}$ is computed from the region marked with the dashed line when $(d_1, d_2)$ is either belong to (0.27 to 0.3,0) or belong to (0, 0.27 to 0.3) of the 2D-FES.

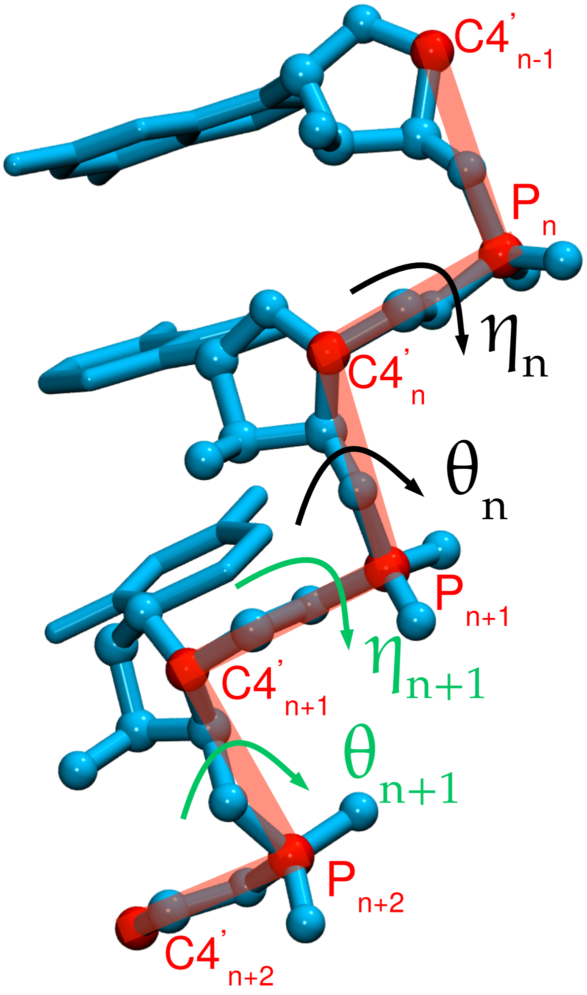


Figure S2: Schematic for $\eta$ and $\theta$ angles for RNA.

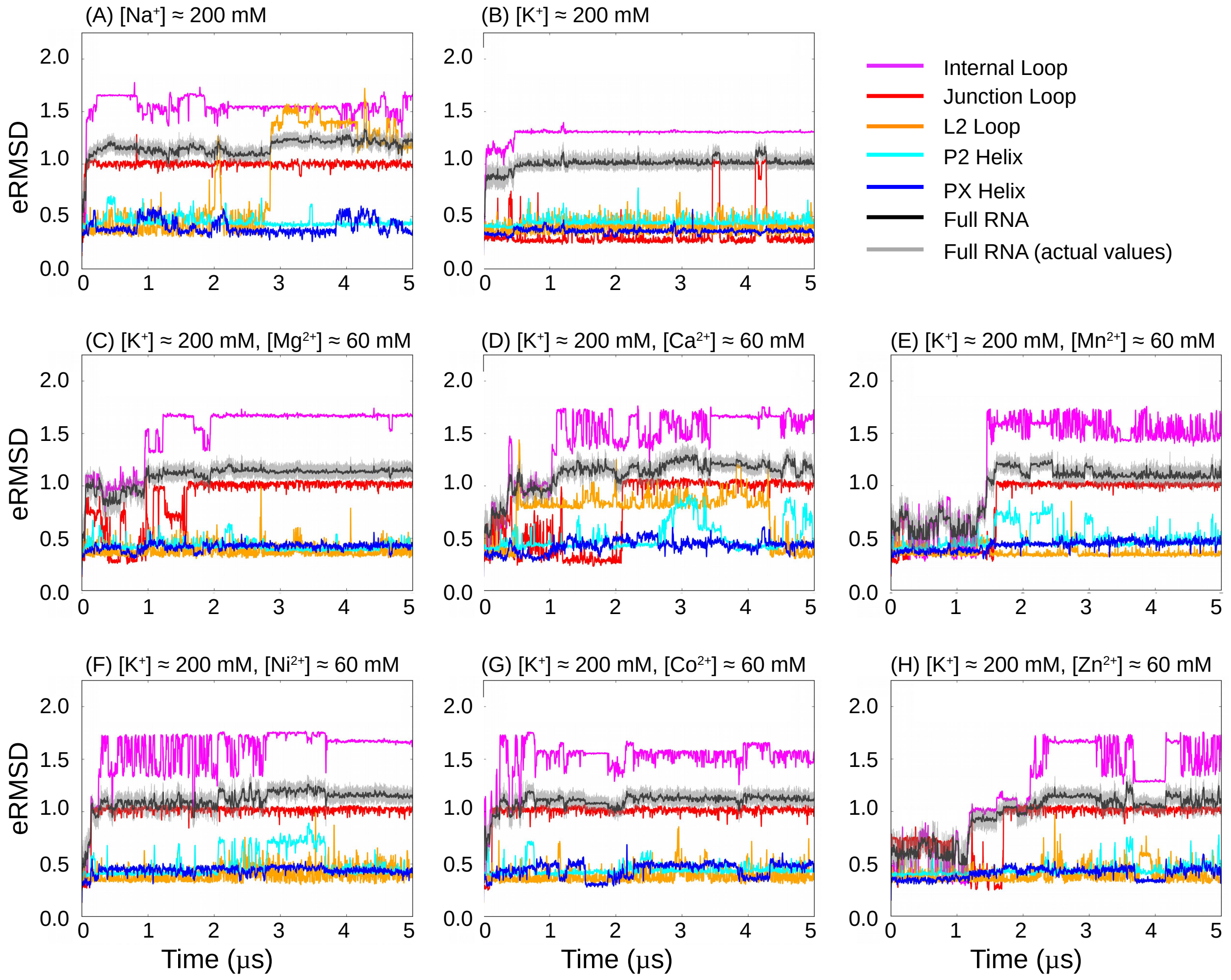


Figure S3: RNA $eRMSD$ with reference to the energy minimized crystal structure is plotted as a function of simulation time. The length of simulation trajectories are $\approx 5\mu s$. Running averages of 5 ns for $eRMSD$ is plotted for the whole RNA and its secondary structural components are shown for the eight different ion solutions: (A) only $Na^+$, (B) only $K^+$, (C) $K^+$ and $Mg^{2+}$, (D) $K^+$ and $Ca^{2+}$, (E) $K^+$ and $Mn^{2+}$, (F) $K^+$ and $Ni^{2+}$, (G) $K^+$ and $Co^{2+}$, and (H) $K^+$ and $Zn^{2+}$. Actual $eRMSD$ values of the whole RNA are shown as gray lines.

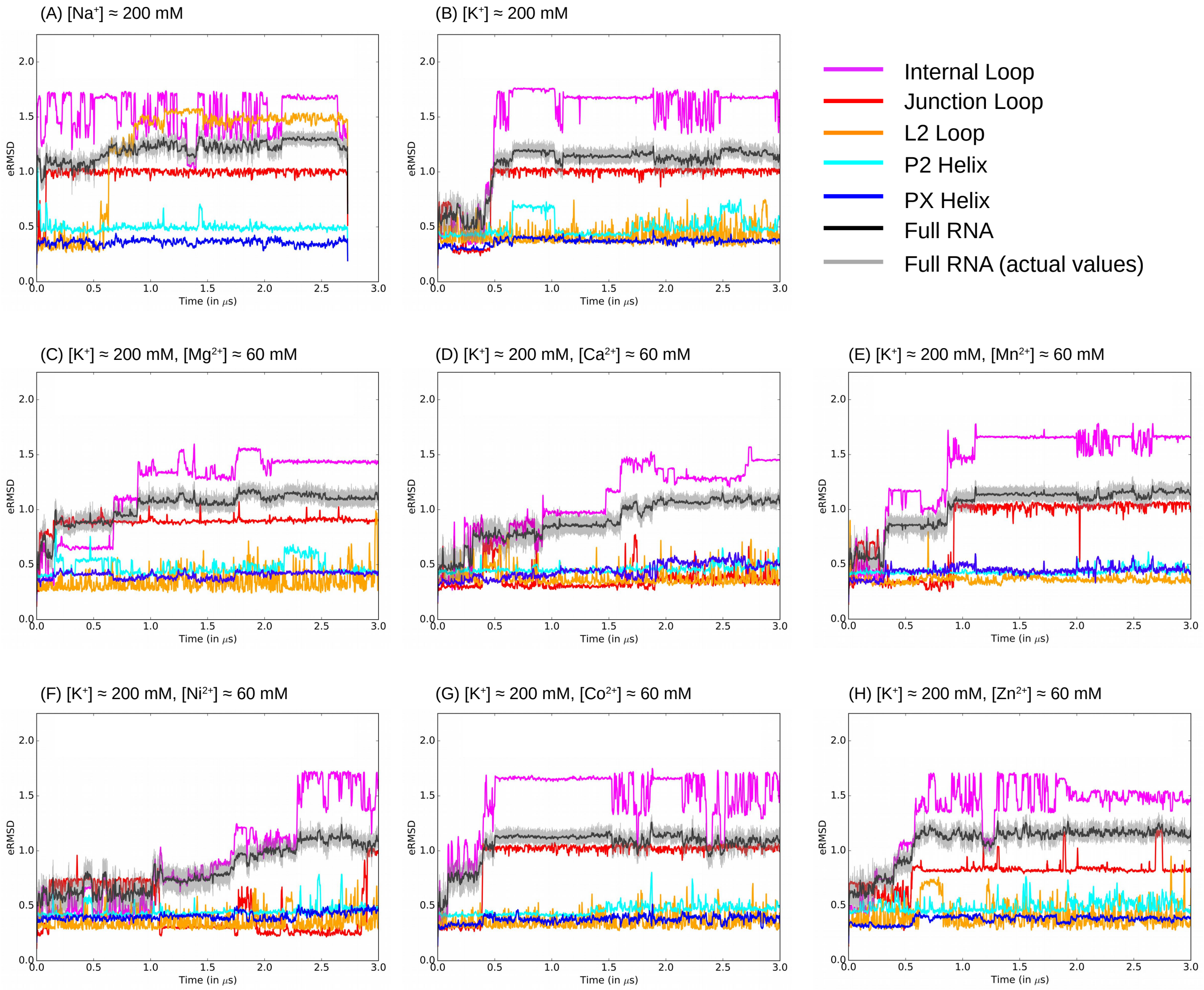


Figure S4: RNA $eRMSD$ with reference to the energy minimized crystal structure is plotted as a function of simulation time for different set of independent simulation trajectories. The length of simulation trajectories are $\approx 3\mu$s. Running averages of 5 ns for $eRMSD$ is plotted for the whole RNA and its secondary structural components are shown for eight different ion solutions: (A) only $Na^+$, (B) only $K^+$, (C) $K^+$ and $Mg^{2+}$, (D) $K^+$ and $Ca^{2+}$, (E) $K^+$ and $Mn^{2+}$, (F) $K^+$ and $Ni^{2+}$, (G) $K^+$ and $Co^{2+}$, and (H) $K^+$ and $Zn^{2+}$. Actual $eRMSD$ values of the whole RNA are shown as gray lines.

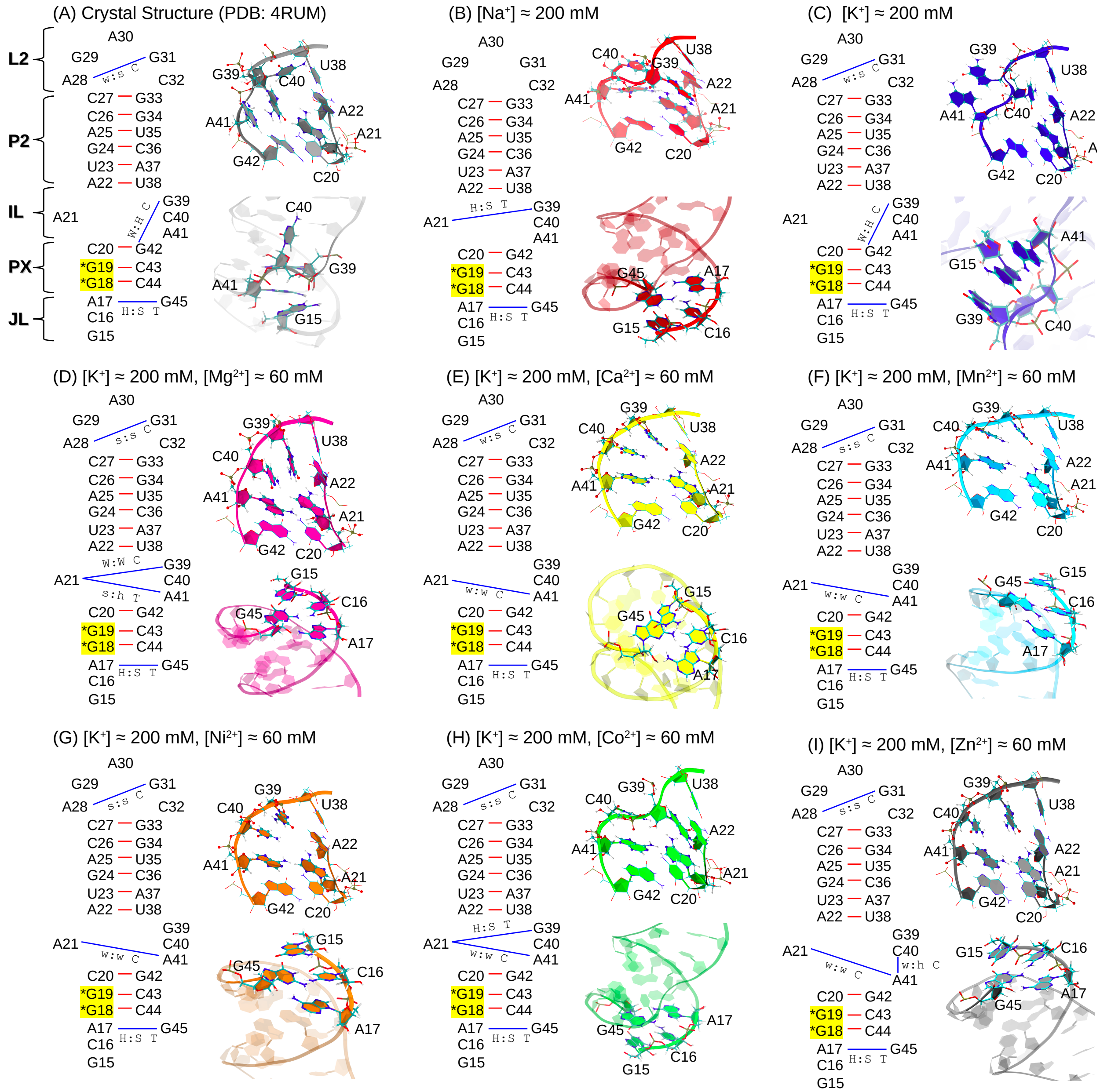


Figure S5: (A) Two-dimensional representation of the native form of the RNA secondary structure. Canonical (red line) and noncanonical (blue line) base-pairing interactions that stabilize the RNA crystal structure are shown schematically. Same analyses are performed on the equilibrated structures observed in the simulations carried out in eight different ion solutions: (B) only $Na^+$, (C) only $K^+$, (D) $K^+$ and $Mg^{2+}$, (E) $K^+$ and $Ca^{2+}$, (F) $K^+$ and $Mn^{2+}$, (G) $K^+$ and $Ni^{2+}$, (H) $K^+$ and $Co^{2+}$, and (I) $K^+$ and $Zn^{2+}$. For better comparison, three-dimensional structures of the internal loop (IL) and junction loop (JL) are shown in each case. The base pairing information is obtained using the BPFIND software (Das et al., J. Biomol. Struct. Dyn., 2006 24, 149-161).

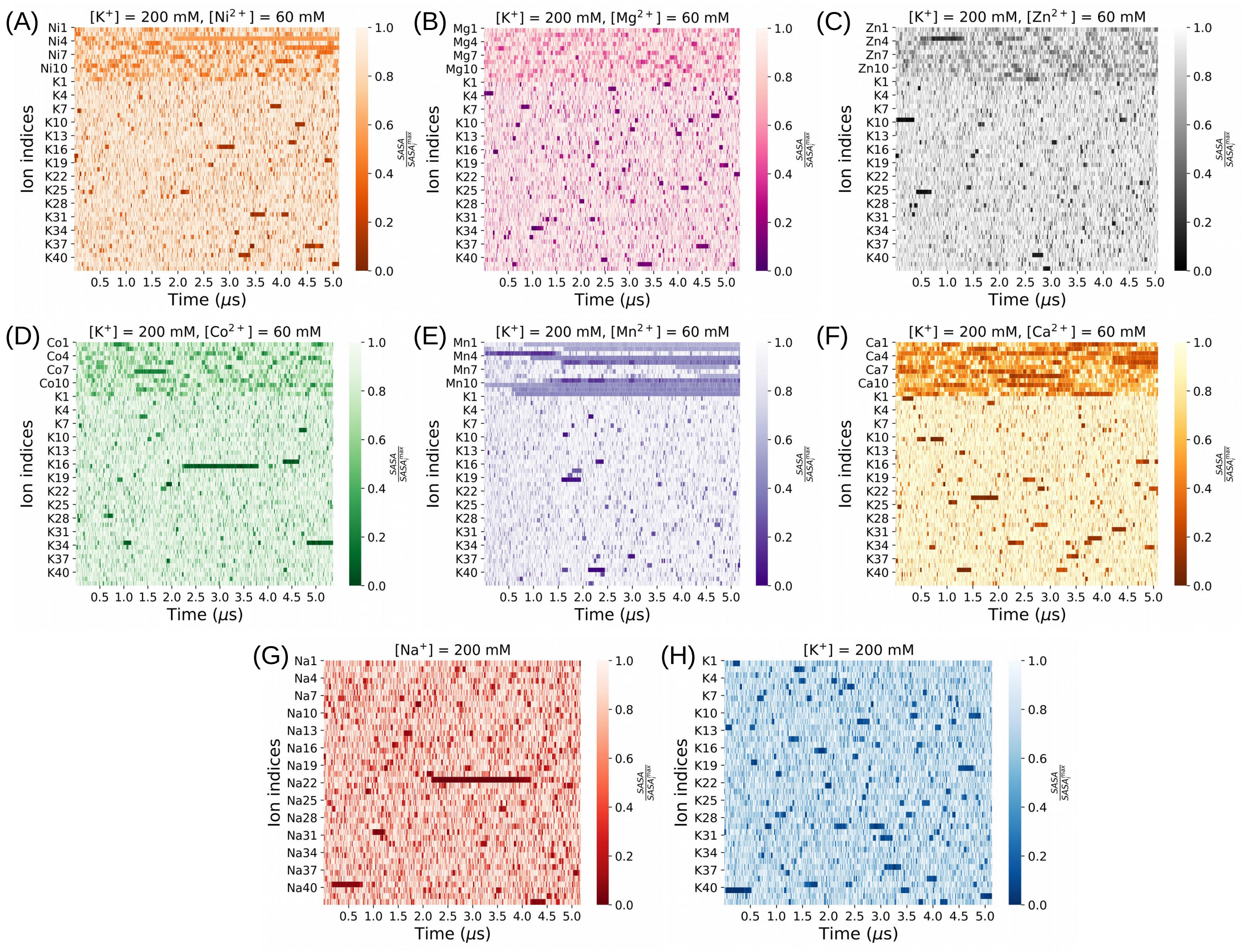


Figure S6: Heatmap of all the individual ions' normalized solvent accessible surface areas ($SASA$). The length of simulation trajectories is $\approx 5\mu$s. The data for eight different ion solutions: (A) only $Na^{+}$, (B) only $K^{+}$, (C) $K^{+}$ and $Mg^{2+}$, (D) $K^{+}$ and $Ca^{2+}$, (E) $K^{+}$ and $Mn^{2+}$, (F) $K^{+}$ and $Ni^{2+}$, (G) $K^{+}$ and $Co^{2+}$, and (H) $K^{+}$ and $Zn^{2+}$.

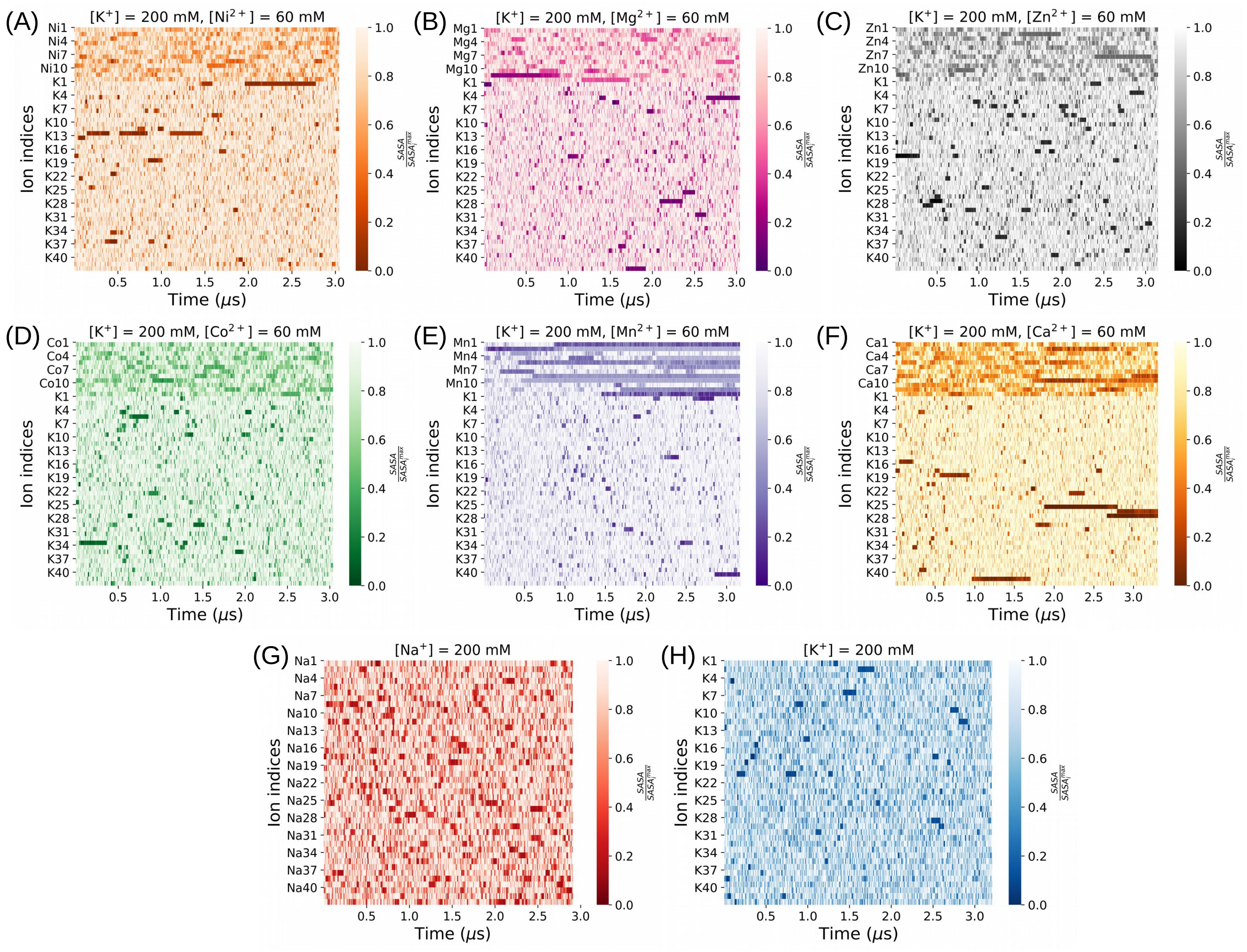


Figure S7: Heatmap of all the individual ions' normalized solvent accessible surface areas ($SASA$) from different set of independent simulation trajectories. The length of simulation trajectories is $\approx 3\mu$s. The data for eight different ions solutions: (A) only $Na^+$, (B) only $K^+$, (C) $K^+$ and $Mg^{2+}$, (D) $K^+$ and $Ca^{2+}$ , (E) $K^+$ and $Mn^{2+}$, (F) $K^+$ and $Ni^{2+}$, (G) $K^+$ and $Co^{2+}$, and (H) $K^+$ and $Zn^{2+}$.

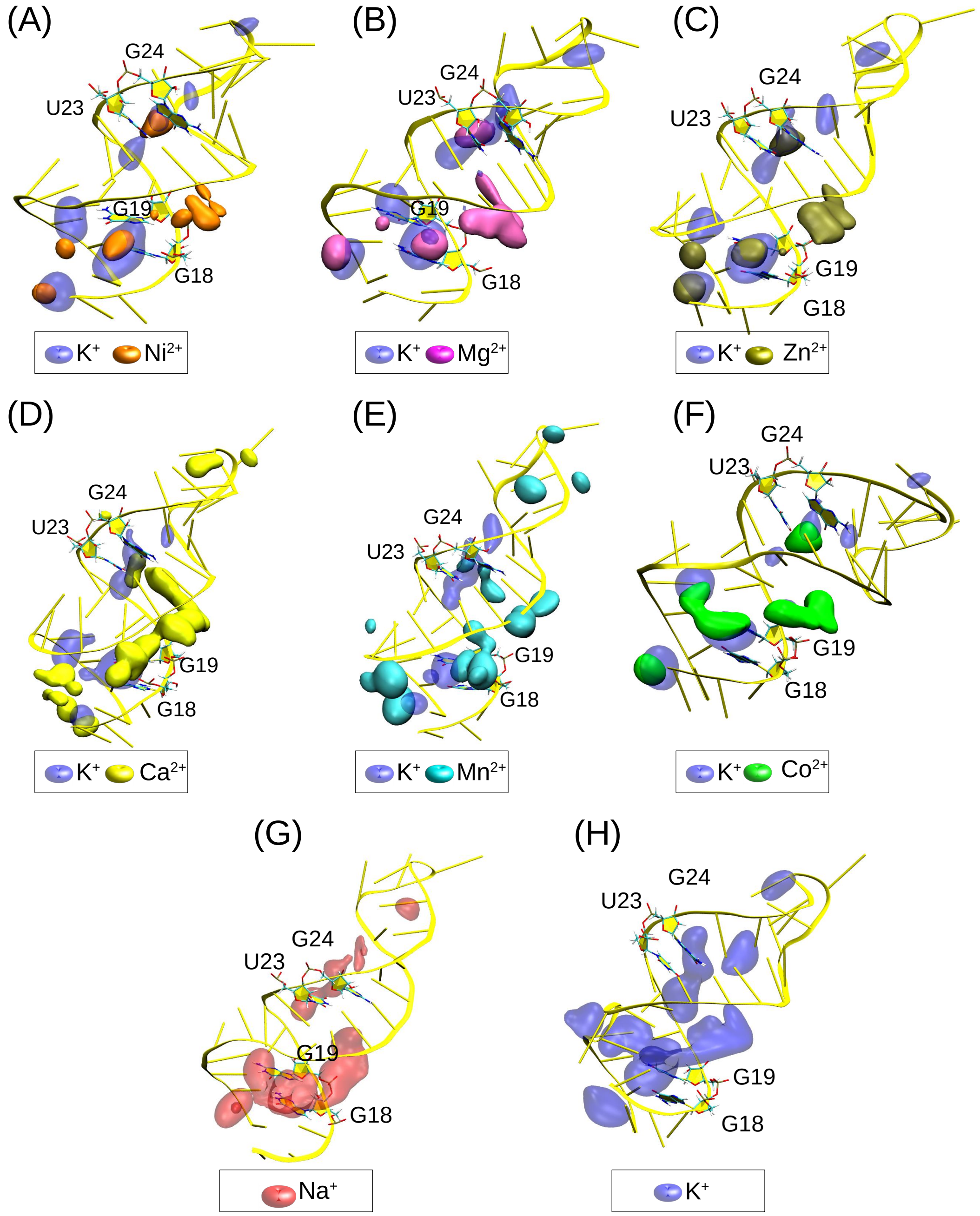


Figure S8: Spatial distribution of mono- and divalent ions around RNA. The isosurfaces corresponding to ISO value 0.03 are plotted for systems (A) $K^+$ and $Ni^{2+}$ (transition metal with high $\rho_c$), (B) $K^+$ and $Mg^{2+}$ (alkaline earth metal with high $\rho_c$), (C) $K^+$ and $Zn^{2+}$ (transition metal with low/high $\rho_c$), (D) $K^+$ and $Ca^{2+}$ (alkaline earth metal with low $\rho_c$), (E) $K^+$ and $Mn^{2+}$ (transition metal with low $\rho_c$), (F) $K^+$ and $Co^{2+}$ (transition metal with high $\rho_c$), (G) $Na^+$, and (H) $K^+$, respectively. The RNA is shown in yellow cartoon representation. Isosurfaces corresponding to $K^+$ are shown in blue in panels (A) to (F). The same for divalent ions are shown using different colors as mentioned in the legend. There are two specific sites on RNA where the ions condense around the nucleobase atoms – (i) around G18 and G19, and (ii) around U23 and G24. These four residues are shown in stick representation using the CPK color scheme.

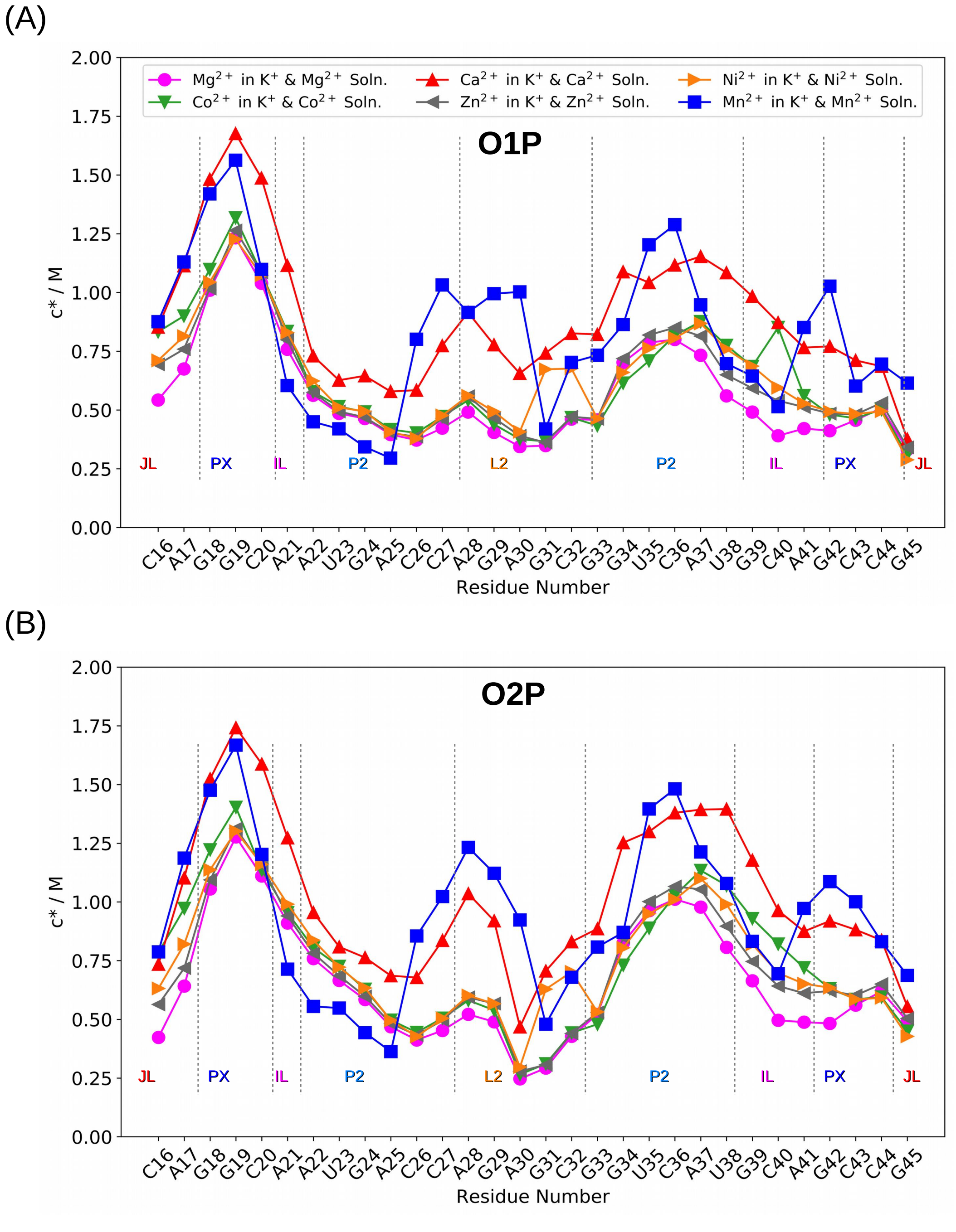


Figure S9: Local ion concentration ($c^*$) around phosphate oxygen atoms (A) O1P and (B) O2P.

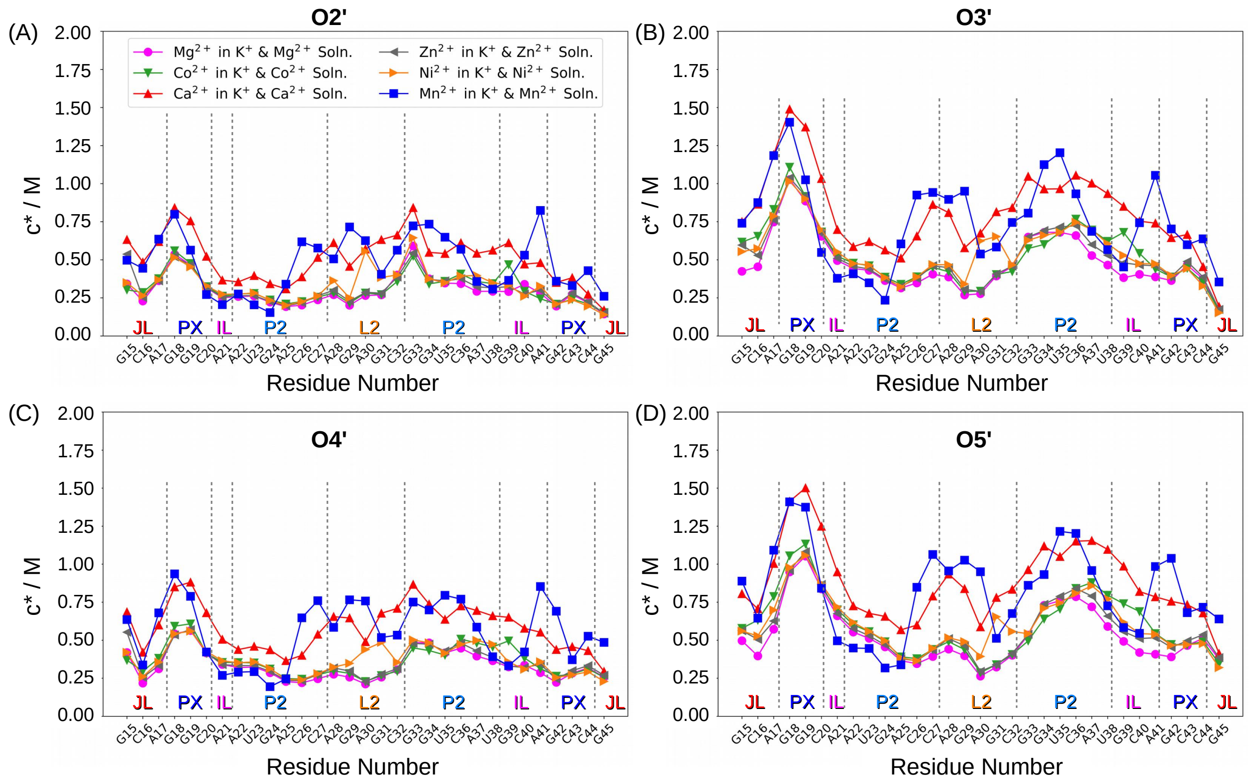


Figure S10: Local ion concentration ($c^*$) around sugar oxygen atoms (A) O2′, (B) O3′, (C) O4′ and (D) O5′. A significantly higher amount of ion condensation is observed around the O atoms, which are involved in the formation of the phosphodiester linkage between two successive nucleotides, i.e., O3′ and O5′. $c^*$ around different nucleobase atoms are illustrated in Figure S20, S21 and S22.

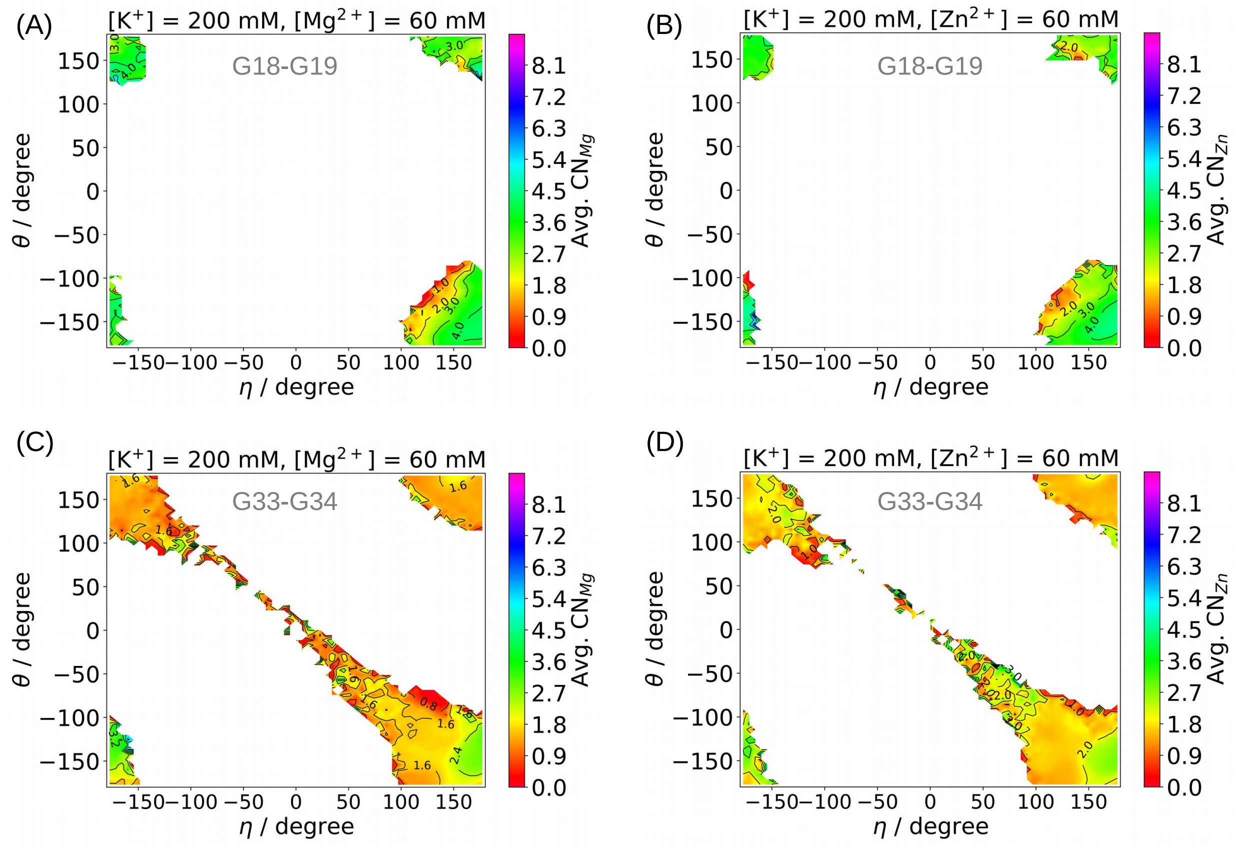


Figure S11: Scatter plots in the $\eta$-$\theta$ space represent the backbone configurations sampled by the G18 and G19 residues during the MD simulations performed in (A) $Mg^{2+}$ and (B) $Zn^{2+}$ solutions. Each point is colored based on the average values of the CN measured between (a) the O atoms attached to the P atom of each residue and (b) all divalent ions present in the solution. The backbone configurations of G18 and G19 residues are compared with the backbone configuration distribution of G33 and G34 residues in (C) $Mg^{2+}$ and (D) $Zn^{2+}$ solutions.

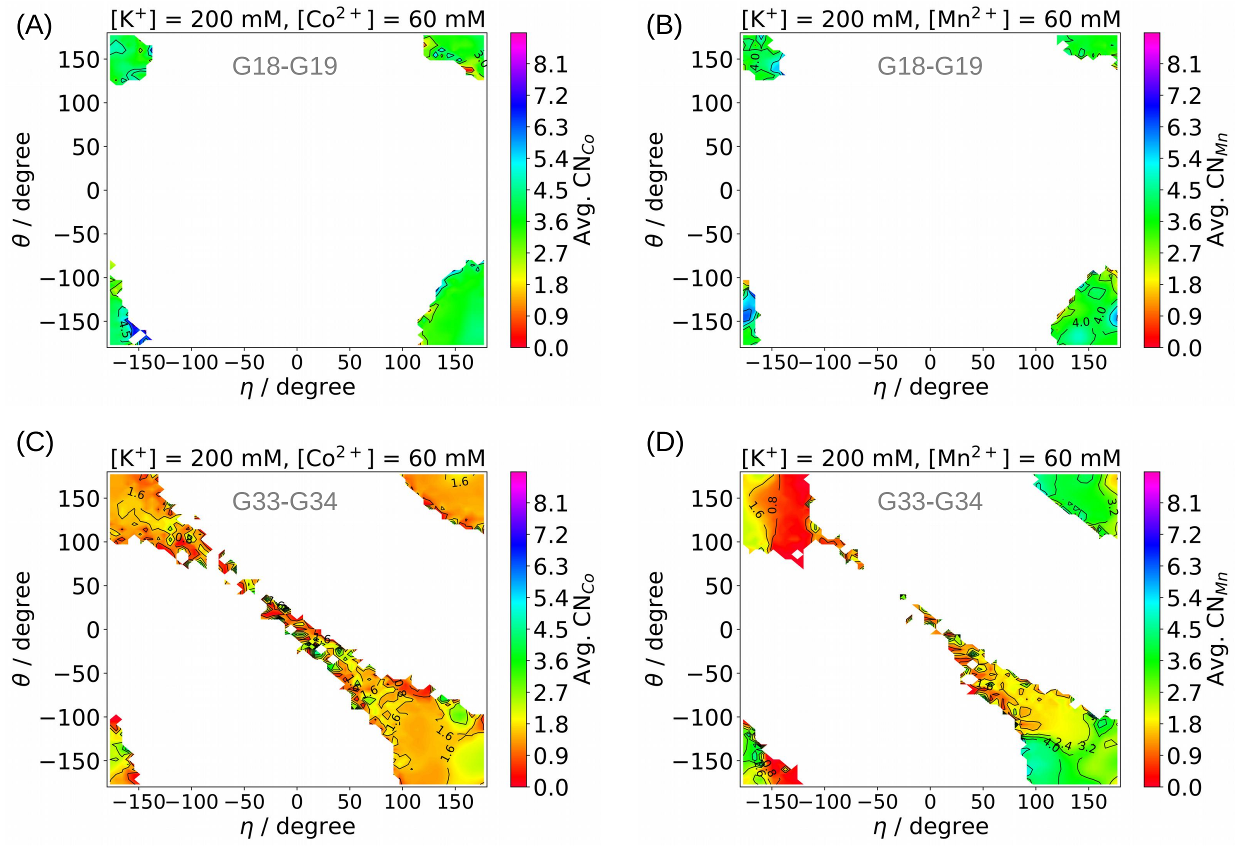


Figure S12: Scatter plots in the $\eta$-$\theta$ space represent the backbone configurations sampled by the G18 and G19 residues during the MD simulations performed in (A) $Co^{2+}$ and (B) $Mn^{2+}$ solutions. Each point is colored based on the average values of the CN measured between (a) the O atoms attached to the P atom of each residue and (b) all divalent ions present in the solution. The backbone configurations of G18 and G19 residues are compared with the backbone configuration distribution of G33 and G34 residues in (C) $Co^{2+}$ and (D) $Mn^{2+}$ solutions.

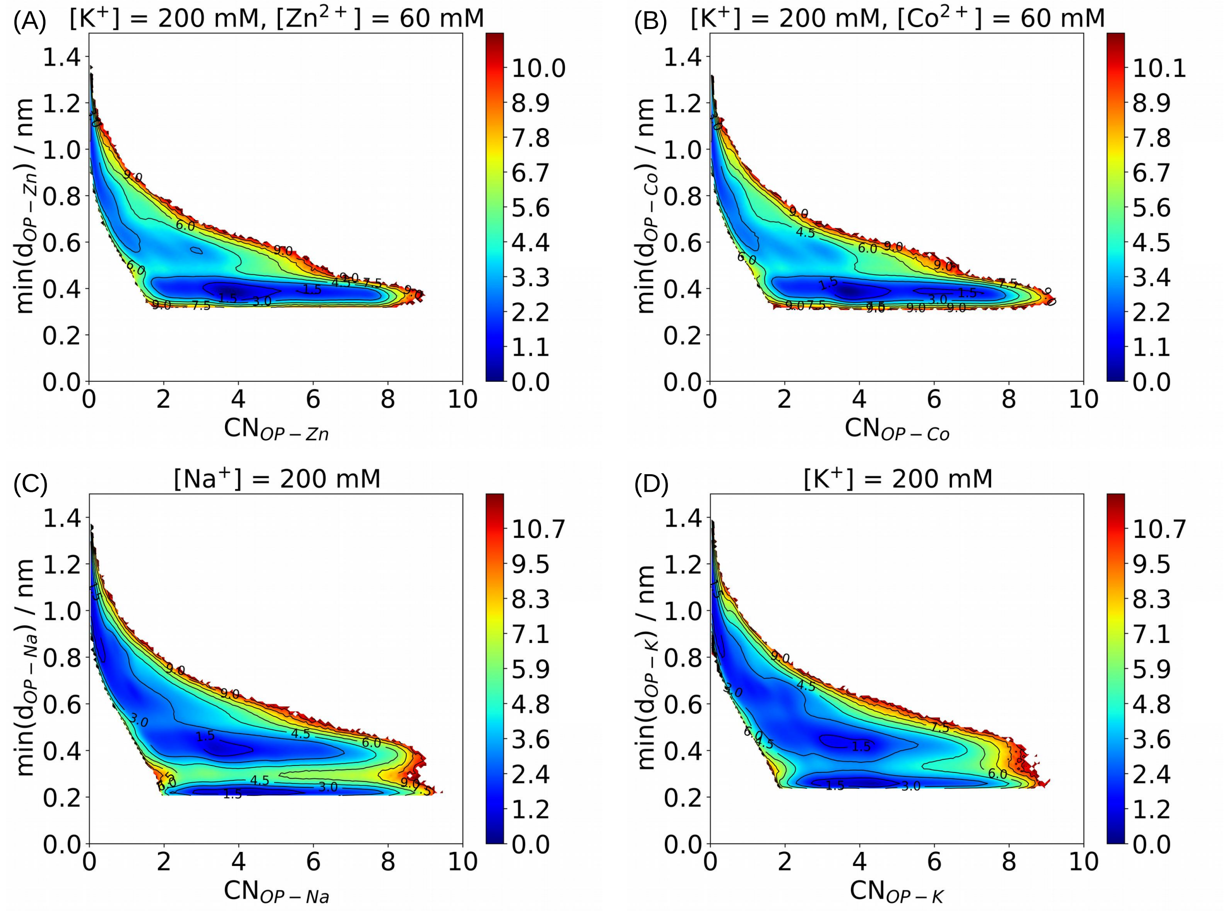


Figure S13: JPD in negative logarithmic scale is plotted for the binding/unbinding of different ions with the RNA backbone for systems (A) $Zn^{2+}$, (B) $Co^{2+}$, (C) only $Na^{+}$, and (D) only $K^{+}$. The horizontal axis represents the coordination number between an individual ion and all the phosphate oxygen atoms. The vertical axis represents the minimum distance between that ion and any phosphate O atoms present in the system.

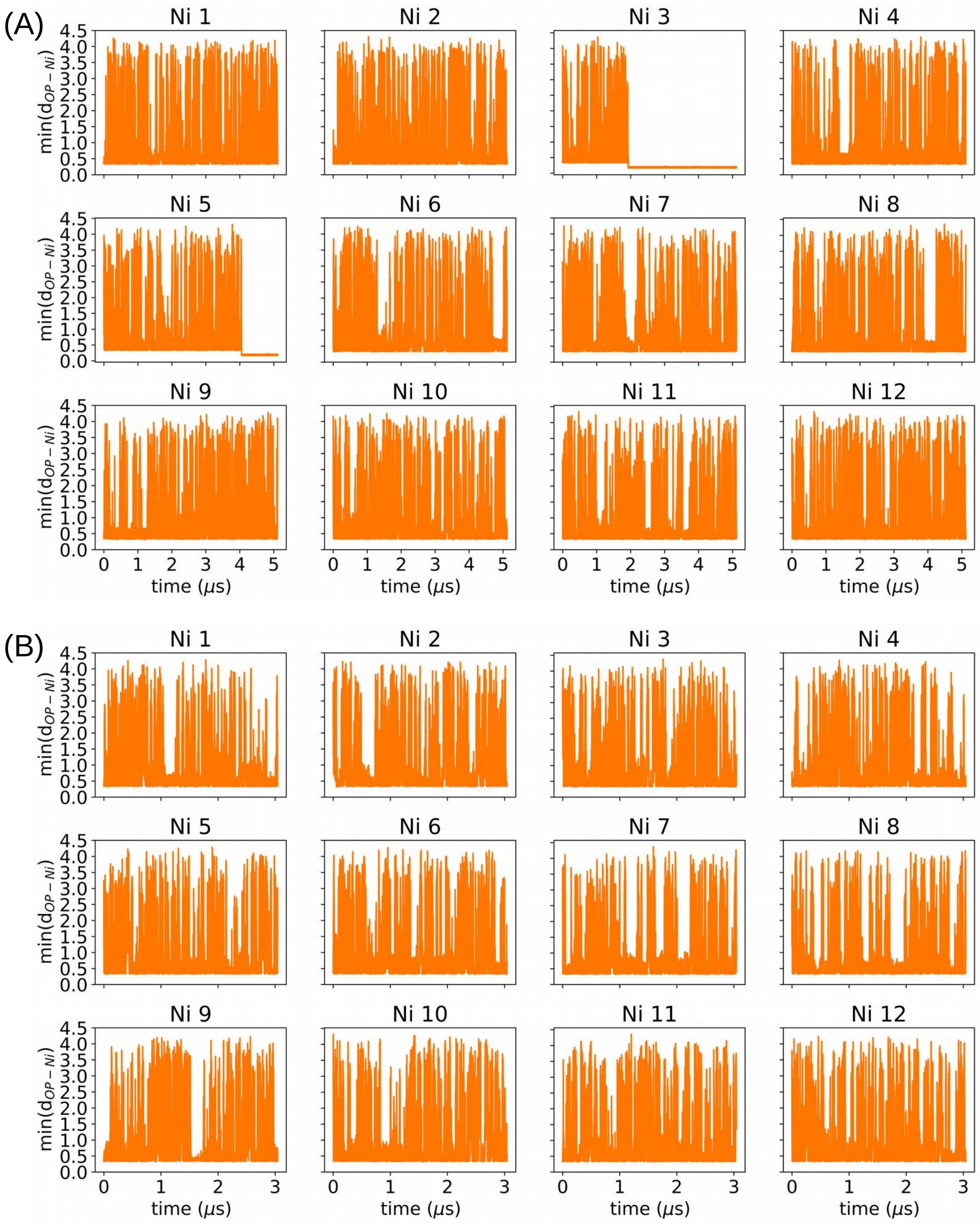


Figure S14: The minimum distance between an individual $Ni^{2+}$ and all the phosphate O atoms (O1P and O2P) in (A) $\approx$ 5 $\mu$s long and (B) $\approx$ 3 $\mu$s long trajectories.

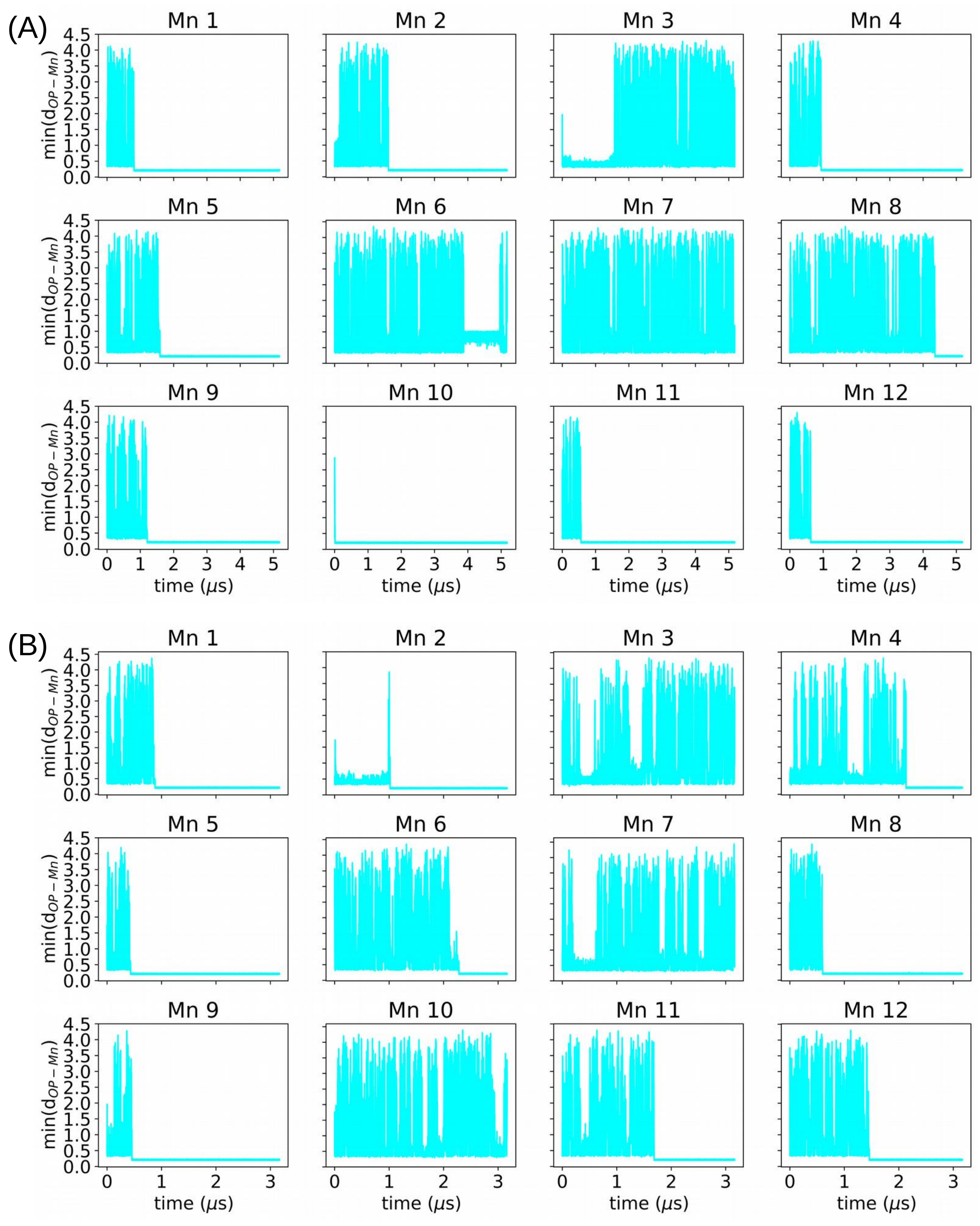


Figure S15: The minimum distance between an individual $Mn^{2+}$ and all the phosphate O atoms (O1P and O2P) in (A) $\approx$ 5 $\mu$s long and (B) $\approx$ 3 $\mu$s long trajectories.

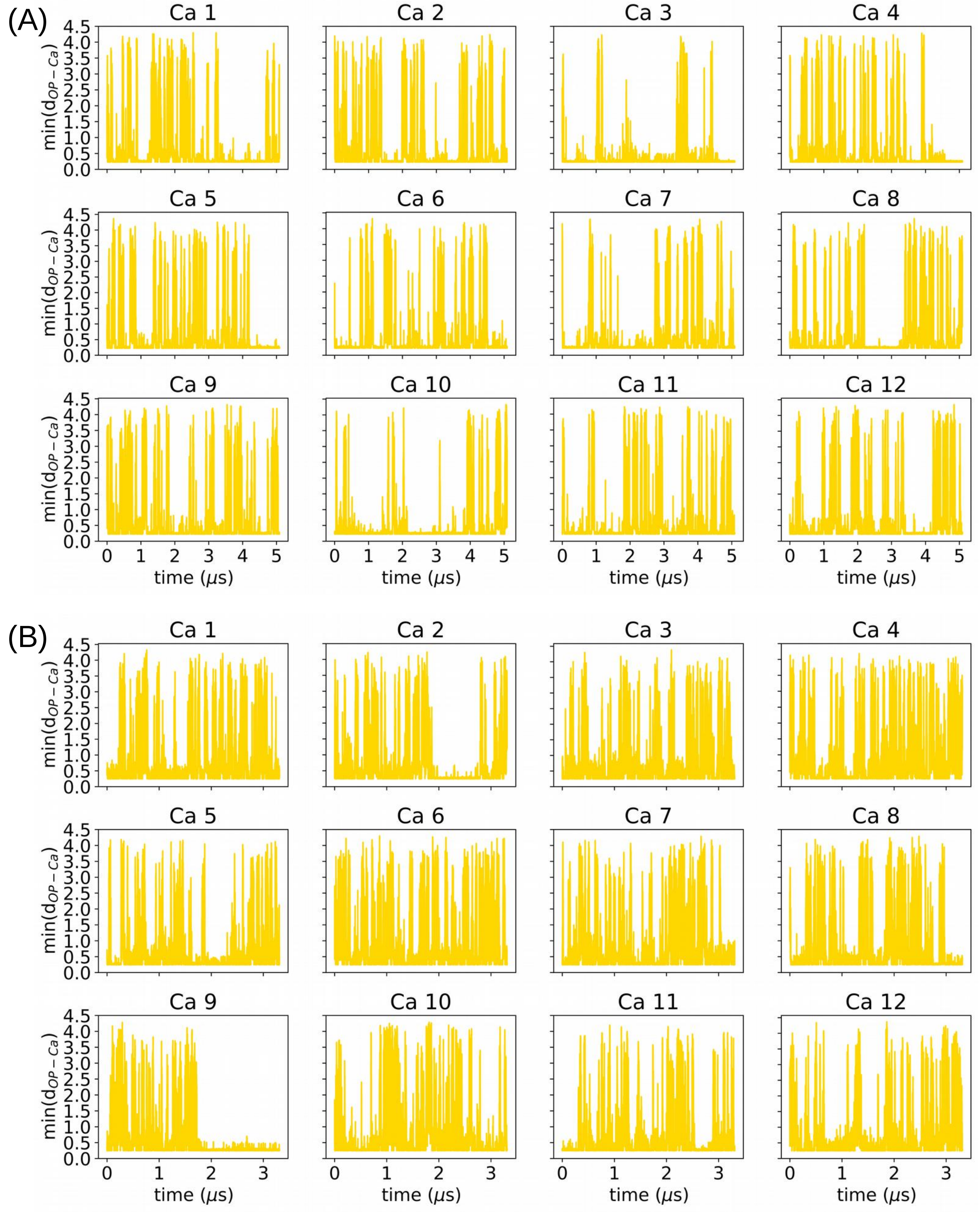


Figure S16: The minimum distance between an individual $Ca^{2+}$ and all the phosphate O atoms (O1P and O2P) in (A) $\approx 5$ $\mu$s long and (B) $\approx 3$ $\mu$s long trajectories.

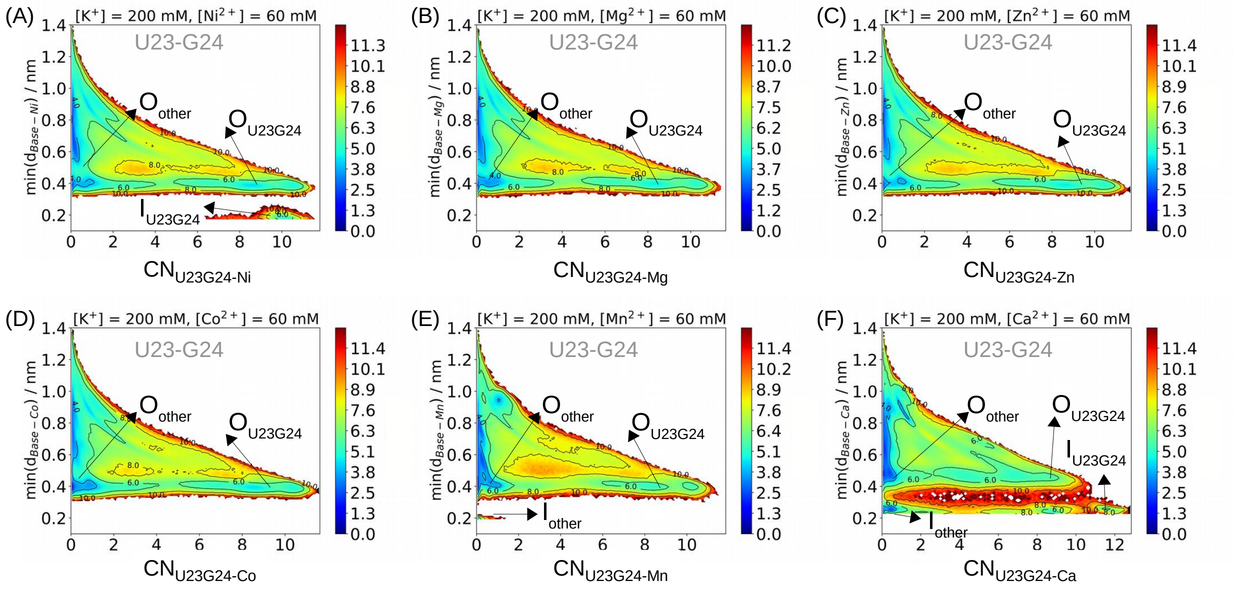


Figure S17: JPD in negative logarithmic scale is plotted for the binding/unbinding of different ions with the Hoogsteen edges of the U23 and G24 residues for the systems (A) $Ni^{2+}$, (B) $Mg^{2+}$, (C) $Zn^{2+}$, (D) $Co^{2+}$, (E) $Mn^{2+}$ and (F) $Ca^{2+}$. The horizontal axis represents the coordination number between divalent ions and selected atoms of the Hoogsteen edges of the nucleobases of U23 and G24. The vertical axis represents the minimum distance between that particular ion and any carbonyl oxygen or imino nitrogen nucleobase atoms present in the system. Energy basins corresponding to the ion's outer shell interaction with the Hoogsteen edge of U23-G24 ($O_{U23G24}$), outer shell interaction with nucleobase atoms other than those of U23-G24 ($O_{other}$), inner shell interaction with the Hoogsteen edge of U23-G24 ($I_{U23G24}$) and inner shell interaction with nucleobase atoms other than those of U23-G24 ($I_{other}$) are labeled.

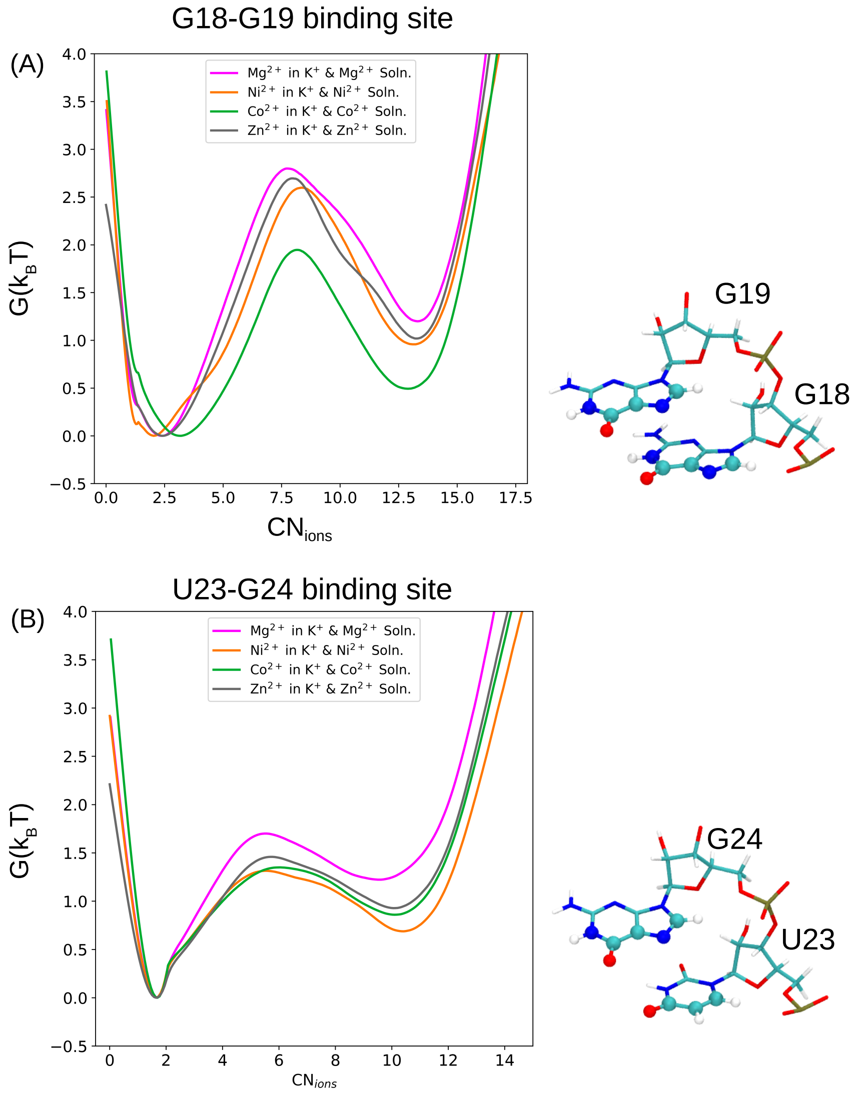


Figure S18: Free energy projected on $CN_{\text{ions}}$ is plotted for the (A) binding of divalent ions at the G18-G19 pocket and (B) binding of divalent ions at the U23-G24 pocket. For each system, $CN_{\text{ions}}$ is calculated between all the divalent ions present in the solution and a set of RNA atoms. The RNA atoms of G18 and G19 that are used in calculating the corresponding $CN_{\text{ions}}$ values are shown as spheres on the right-hand side of panel-A. Similarly, the RNA atoms of U23 and G24 that were used in calculating the corresponding $CN_{\text{ions}}$ values are shown as spheres on the right-hand side of panel-B.

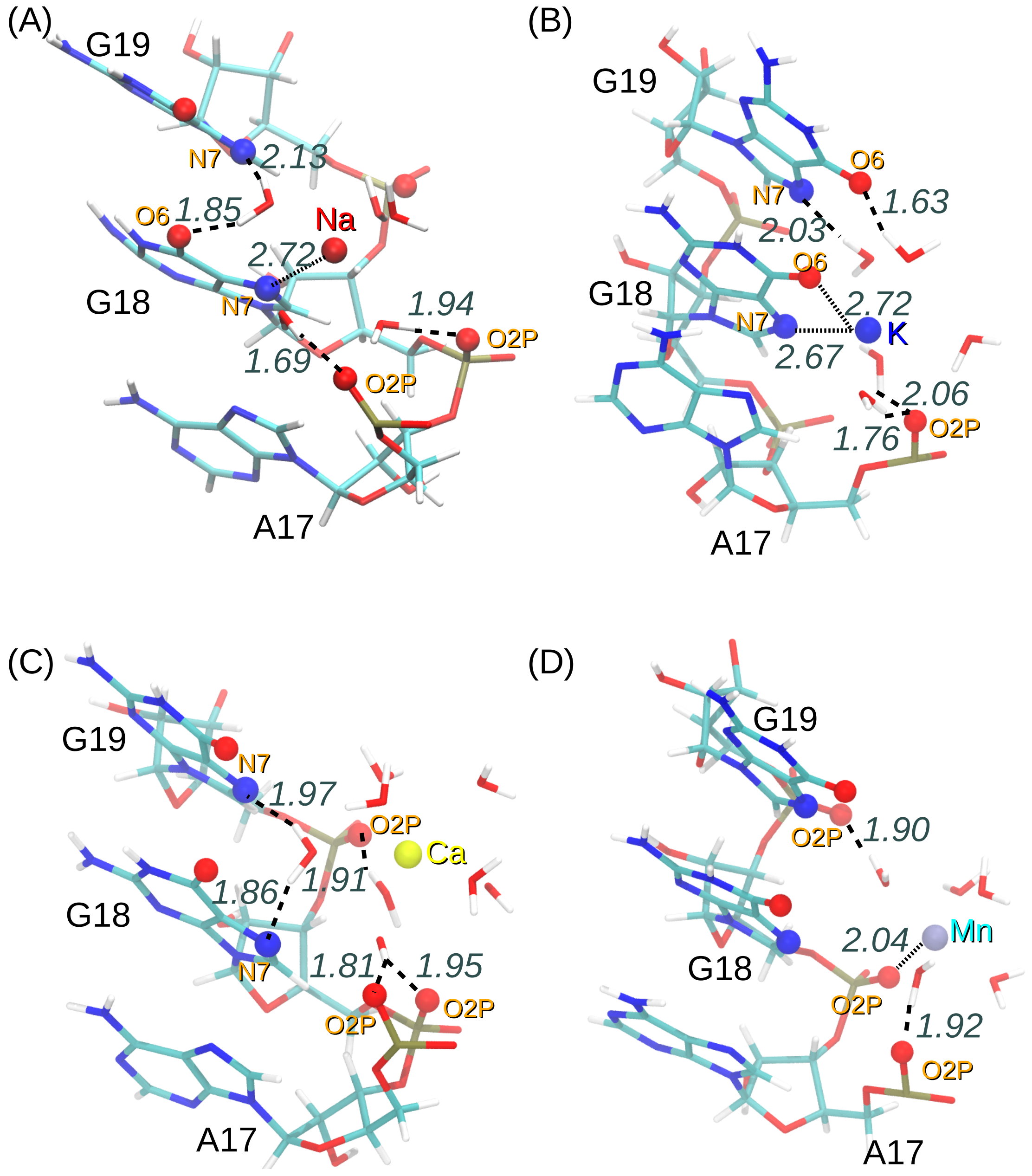


Figure S19: Ion-RNA interaction at the G18-G19 site is shown for (A) $Na^{+}$, (B) $K^{+}$, (C) $Ca^{2+}$, and (D) $Mn^{2+}$ ions. The bound ion is shown as a sphere, and its coordinated water molecules are shown in stick representation. Hydrogen bonds between the ion coordinated water molecules and RNA atoms are shown as black broken lines, and the corresponding hydrogen-acceptor distances are reported in Å.

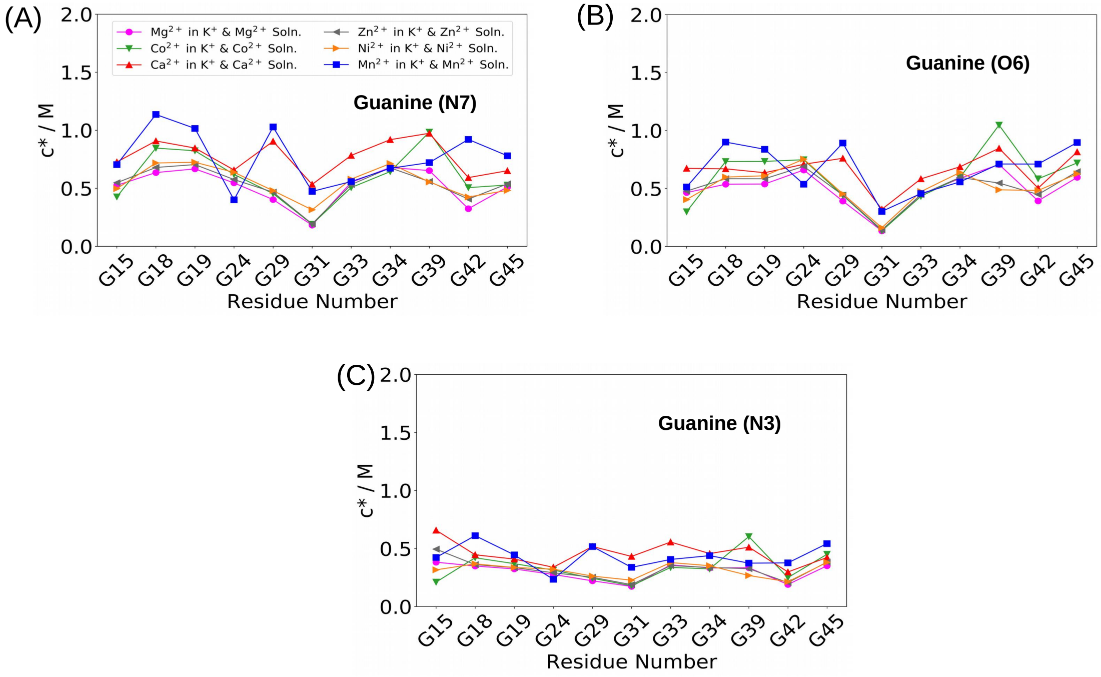


Figure S20: Local ion concentration ($c^*$) around the (A) N7, (B) O6, and (C) N3 atoms in the guanine residues (G15, G18, G19, G24, G29, G31, G33, G34, G39, G42, and G45) present in the RNA structure.

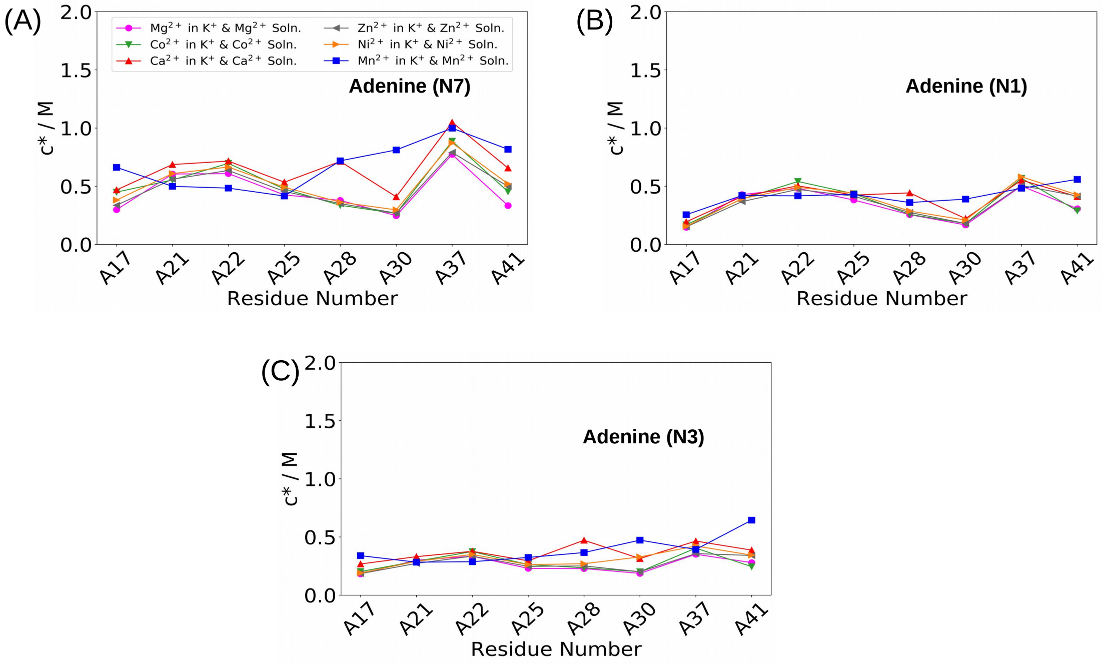


Figure S21: Local ion concentration ($c^*$) around the (A) N7, (B) N1, and (C) N3 atoms in the adenine residues (A17, A21, A22, A25, A28, A30, A37 and A41) present in the RNA structure.

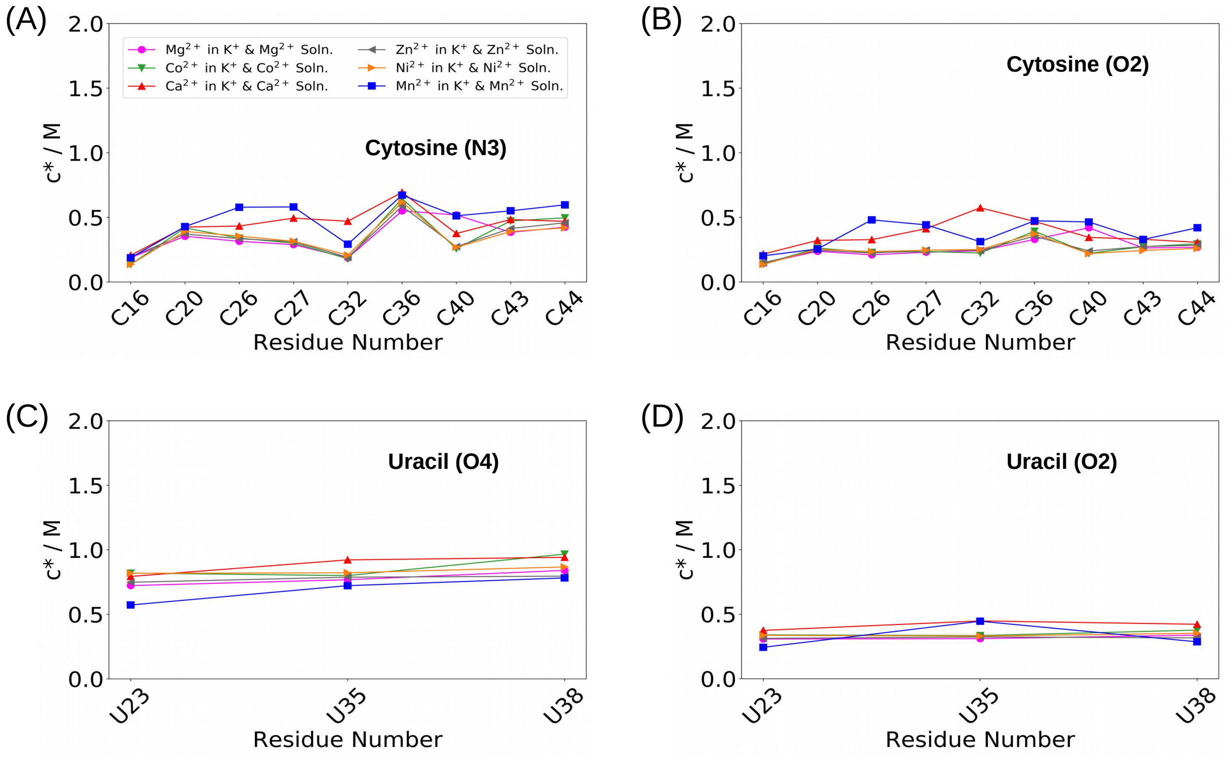


Figure S22: Local ion concentration ($c^*$) around the imino nitrogen and carbonyl oxygen atoms of different nucleobases: (A) N3 of cytosine, (B) O2 of cytosine, (C) O4 of uracil, and (D) O2 of uracil present in the RNA structure.
